# Platform-Imprinted Transcriptional and Clonal Remodeling of αβ and γδT Cells After Allogeneic Transplantation

**DOI:** 10.64898/2026.02.09.704768

**Authors:** A.H.G. Stuut, P. Brazda, A. Janssen, A. Vyborova, E. Karaiskaki, F. Keramati, D.A. de Bont, M.J.T. Nicolasen, L.C.D.E. Gatti, T.J.A. Hutten, J. Yildiz, H.T. Spierings, G.C.M. Straetemans, D.X. Beringer, S. Pagliuca, H.G. Stunnenberg, Z. Sebesyten, J. Drylewicz, M.A. de Witte, J. Kuball

## Abstract

Immune reconstitution after allogeneic hematopoietic stem cell transplantation is influenced by graft-composition and viral reactivation, but the combined long-term impact on αβ and γδT cells remains unclear. We analyzed a cohort of 213 patients receiving either αβT cell-depleted grafts (n=146; graft engineering that removes donor αβT cells) or T cell-replete grafts (n=67; containing donor T cells). Longitudinal immune phenotyping was integrated with bulk and single-cell TCR repertoire and transcriptomic profiling. CMV reactivation was associated with expansion of CD8^+^ αβT cells across both transplant types and with numerical dominance of Vδ2^−^ γδT cells specifically in αβT cell-depleted recipients. Vδ2^−^ γδT cells underwent early polyclonal expansion followed by repertoire focusing, independent of CMV, whereas αβT cells remained clonally restricted. Reduced early Vδ2^+^ γδTCR diversity was associated with EBV reactivation. Single-cell and TCR tracking analyses revealed long-term persistence of donor-derived Vδ2^+^ γδTCRs, whereas Vδ1^+^ γδ and αβT cell repertoires were predominantly rebuilt de novo. Despite de novo rebuilding, αβTCR repertoire diversity diverged by platform at one year: αβT cell-depleted recipients exhibited marked (hyper)expansion of αβTCR clonotypes and lower diversity than T cell-replete recipients, indicating a durable imprint of graft engineering on αβTCR-clonality. Transcriptomic profiling showed that post-transplant T cells predominantly adopted effector programs, with platform-dependent polarization toward cytotoxic signatures in αβT cell–depleted recipients and toward AREG-associated tissue-repair signatures in T cell–replete recipients, consistent with wound-healing functions. In conclusion, transplantation platforms imprint durable clonal and transcriptional remodeling of αβ and γδT cells, while viral reactivation primarily amplifies expansion without fundamentally reshaping repertoire architecture.

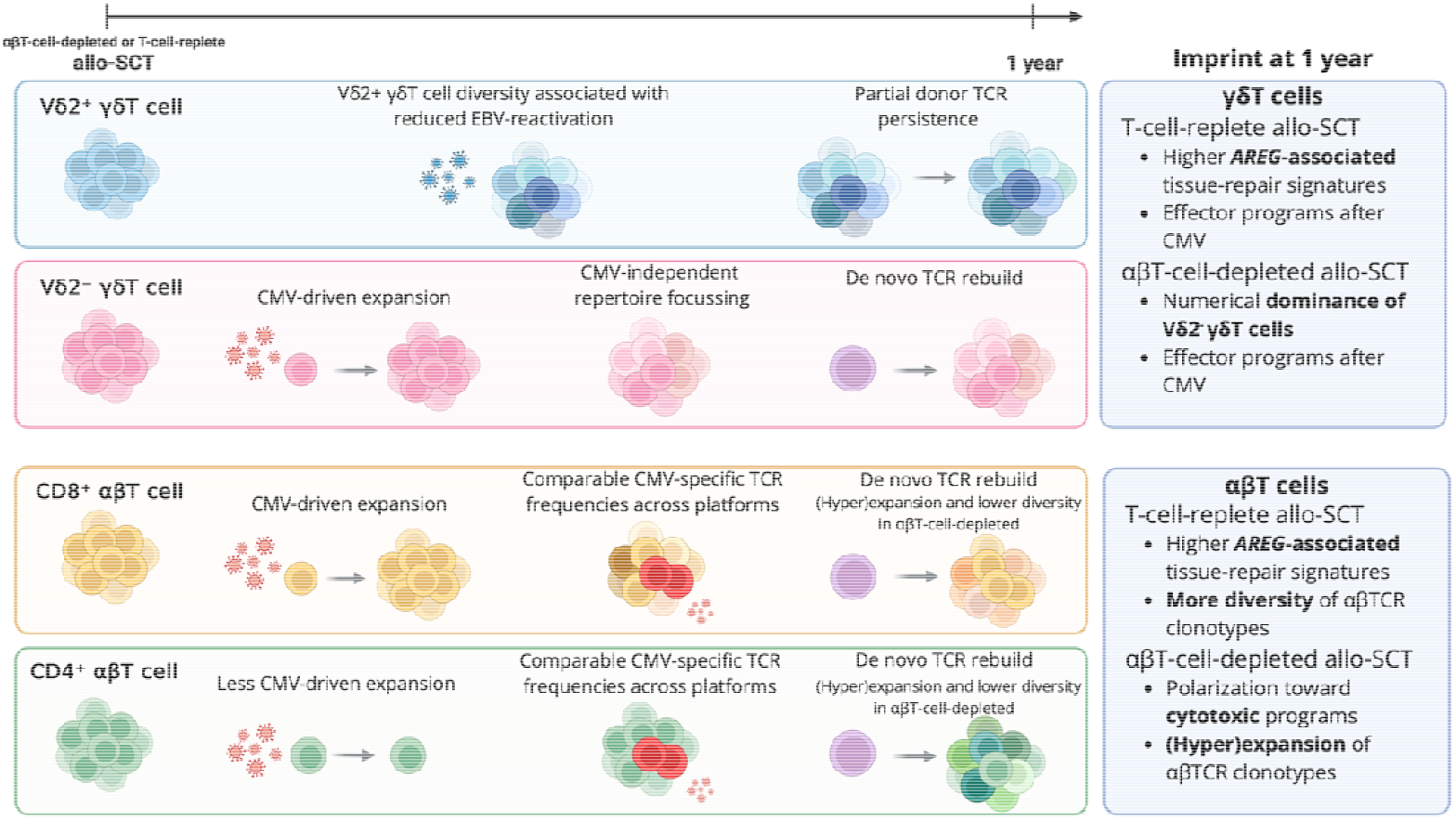

## Introduction

A well-balanced immune reconstitution is critical for the success of allogeneic hematopoietic stem cell transplantation (allo-HSCT), as it determines a patient’s ability to control complications such as graft-versus-host disease (GVHD) and viral infections. However, immune reconstitution dynamics vary substantially across transplantation platforms, with pronounced differences between αβT cell-depleted and T cell-replete grafts^1-6^. αβT cell-depleted grafts reduce transfer of mature donor αβ T cells to mitigate GVHD but may delay adaptive T cell recovery, while T cell-replete grafts provide immediate donor T cell immunity but carry a higher risk of alloreactivity^1^. These platform-specific differences affect the timing and composition of immune subset recovery and, consequently, host responses to viral challenges such as cytomegalovirus (CMV) and Epstein–Barr virus (EBV), both of which frequently reactivate post-transplantation and remain major contributors to morbidity and mortality ^7,8^.

Most studies examining the interplay between immune recovery and viral reactivation have been conducted in T cell-replete settings. In this context, early reconstitution of NK cells and CD4^+^ αβ T cells has been associated with protection against CMV reactivation, but not EBV reactivation^9^. CMV reactivation has been shown to drive numerical expansion and effector differentiation of Vδ2^−^ γδT cells and CD8^+^ αβT cells^9-14^, accompanied by clonal proliferation and reduced TCR repertoire diversity, as revealed by next-generation sequencing studies^15-19^. In contrast, EBV reactivation typically occurs in the setting of insufficient immune control. It does not promote immune recovery, which has motivated the development of EBV-specific adoptive T cell therapies to prevent post-transplant lymphoproliferative disease^20^. However, as transplantation platforms evolve, these established immune–virus relationships may no longer directly translate.

This has become increasingly relevant with the growing clinical implementation of ex vivo αβT cell-depleted grafts^21,22^, which introduce fundamentally distinct immune reconstitution dynamics^1,2^. Profound delay of αβT cell recovery shifts early immune responses toward innate and unconventional lymphocyte populations, including NK and γδT cells^2,4,6,23-25^. In this αβT cell-depleted setting, viral reactivations are likely to shape immune phenotypes differently than in T cell-replete platforms. For instance, CMV reactivation in αβT cell-depleted or umbilical cord blood recipients has been associated with the expansion of memory-like CD57^+^NKG2C^+^ NK cells^26-29^. Despite these emerging observations, systematic comparative analyses across platforms and immune subsets remain limited.

To address this gap, we investigated how αβ T cell–depleted versus T cell–replete transplantation platforms shape post-transplant immune reconstitution, and how CMV and EBV reactivations modulate this process. We performed longitudinal immune profiling and integrated bulk and single-cell TCR repertoire and transcriptomic analyses to characterize reconstitution of both αβ and γδT cell compartments across platforms. In addition, we performed single-cell RNA sequencing on donor samples and on recipient samples one year post-transplantation from both platforms in patients with CMV reactivation to characterize phenotypic and transcriptional changes in T cell subsets. We demonstrate that the transplantation platform establishes the baseline long-term framework of post-transplant immunity. At the same time, CMV reactivation amplifies this framework by driving distinct transcriptional and functional polarization of expanding CD8^+^ αβ and Vδ2^−^ γδT cell subsets.

## Materials & Methods

### Clinical cohort

Patients with hematological malignancies who received an αβT cell-depleted stem cell transplantation at the University Medical Center Utrecht (UMCU) between 2015 and 2017 were included in this retrospective study. Conditioning consisted of anti-thymocyte globulin (ATG), fludarabine, and busulfan. αβ T cell depletion was performed using anti-αβTCR antibodies and the CliniMACS system, allowing a maximum residual αβ T cell dose of 5 × 10□ cells/kg. Mycophenolate mofetil (MMF) was used as single GVHD prophylaxis, and pre-emptive donor lymphocyte infusion (DLI; 1×10□ CD3^+^ T cells/kg) was administered at three months post-transplantation in eligible patients. As a comparator group, T cell-replete transplant recipients treated at UMCU between 2011 and 2018 were included after either reduced-intensity or myeloablative conditioning with standard GVHD prophylaxis. Peripheral blood stem cells were used in both cohorts. Immune reconstitution and viral reactivations were monitored longitudinally as part of routine care, with immune subset recovery assessed by flow cytometry. CMV reactivation was defined as >250 IU/mL, and EBV reactivation as >1000 IU/mL by PCR. All patients provided written informed consent, and the study was approved by the local ethics committee (METC nr 21-322, UMCU) (see^30,31^ and **Supplementary Material**).

### T cell receptor repertoire sequencing and single-cell profiling

To characterize the clonal architecture and transcriptional programs of αβ and γδ T cell subsets during post-transplant immune reconstitution, we combined bulk next-generation sequencing of the T cell receptor (TCR) with paired single-cell RNA sequencing (scRNA-seq) and single-cell TCR sequencing (scTCR-seq). These complementary approaches enabled quantitative assessment of repertoire diversity, clonal expansion, and functional polarization at both population and single-cell resolution. TCRβ and TCRδ repertoires were sequenced from flow-sorted CD4^+^ αβ, CD8^+^ αβ, and γδ T cell subsets using established amplification protocols^32^, followed by Illumina-based sequencing. Productive CDR3 sequences were extracted using MiXCR^33^, normalized for sequencing depth, and analyzed with the immunarch package^34^. For single-cell profiling, αβ and γδ T cells were flow-sorted from PBMCs collected approximately one year post-transplantation and processed using the 10x Genomics Chromium 5′ platform with paired V(D)J enrichment, including custom γδTCR primers. scRNA-seq data were processed with Cell Ranger and analyzed in Seurat^35-38^, with reference-based cell-type annotation using Azimuth^37,39^. Furthermore, we used the T cellAnnoTator (TCAT) pipeline to characterize functional programs in T cells^40^. αβTCRs were screened for CMV specificity using the scRepertoire and Trex packages^41,42^, which annotate αβTCRs with epitope data from VDJdb, McPAS-TCR, IEDB, and PIRD databases^43-46^. Complete experimental protocols and detailed analytical workflows for TCR sequencing, scRNA-seq, and scTCR-seq are provided in the **Supplementary Material**.

## Results

### CMV-driven divergence in long-term T cell recovery across transplantation platforms

To compare immune reconstitution following αβT cell-depleted-(n = 146) versus T cell-replete allo-HSCT (n = 67) (baseline characteristics in **Table 1**), we longitudinally assessed the numerical recovery of CD8^+^ αβT cells, CD4^+^ αβT cells, Vδ2^+^ γδT cells, and Vδ2^−^ γδT cells over the first three years post-transplantation. Given the known impact of CMV reactivation on post-transplant immunity^11,15^, patients were stratified by CMV reactivation status. CMV reactivation occurred at 27 days (median, range 0– 202) after αβT cell-depleted and 33 days (median, range 3–237) after T cell-replete transplantation. In patients without CMV reactivation, CD4^+^ and CD8^+^ αβT cells reconstituted more rapidly and reached higher absolute counts after T cell-replete transplantation (**Figure 1A, B**). In CMV-reactivated patients, a distinct numerical recovery pattern emerged. CD8^+^ αβT cells expanded markedly in both transplant platforms (**Figure 1C,D; Supplementary Figure S1**). In contrast, CD4^+^ αβT cell recovery was slow and was not augmented by CMV reactivation (**Figure 1C,D**). Across both platforms, the Vδ2^+^/Vδ2^−^ γδT cell ratio declined over time, with a sustained inversion in patients who received an αβT cell-depleted allograft and/or experienced CMV reactivation (**Supplementary Table S1**). Notably, from the earliest phase of immune reconstitution (median day 23), CMV reactivation in the αβT cell-depleted platform was associated with a sustained numerical expansion of Vδ2^−^ γδT cells, which resulted in an inversion of the Vδ2^+^/Vδ2^−^ γδT cell ratio that persisted for up to three years (**Figure 1C; Supplementary Figure S1A**). Alongside CD8^+^ αβT and Vδ2^−^ γδT cells, NK cells also expanded in association with CMV reactivation. This effect was significant in the αβT cell-depleted platform (p = 0.010; **Supplementary Figure S1A**) and similarly observed in the T cell-replete platform (**Supplementary Figure S1B**). B cell numbers were unaffected by CMV reactivation in both platforms. Together, these data demonstrate that while CMV reactivation serves as a strong numerical amplifier of CD8^+^ αβT cells across transplant platforms, αβT cell depletion uniquely imprints a long-lasting inversion of the Vδ2^+^/Vδ2^−^ γδT cell balance.

**Table 1.** Baseline characteristics.

|  | **Ex vivo αβT-cell**<br>**depleted**<br>N = 146 | **T-cell-replete**<br>N = 67 | **p-value** |
| --- | --- | --- | --- |
| **Median age (IQR)** | 53 (43, 64) | 53 (45, 62) | 0.84<sup>1</sup> |
| **Male, n (%)** | 95 (65%) | 47 (70%) | 0.47<sup>2</sup> |
| **Median follow-up post allo-HSCT in**<br>**months (IQR)** | 35 (6, 60) | 60 (27, 100) | <0.001<sup>1</sup> |
| **Year of SCT, n (%)** |  |  | <0.001<sup>2</sup> |
| 2011-2015 | 53 (36%) | 55 (82%) |  |
| 2016-2022 | 93 (64%) | 12 (18%) |  |
| **Disease, n (%)** |  |  | 0.22<sup>2</sup> |
| ALL | 21 (14%) | 8 (12%) |  |
| AML/MDS RAEB2 | 48 (33%) | 20 (30%) |  |
| LYMPHOMA | 20 (14%) | 19 (28%) |  |
| MDS | 14 (10%) | 6 (9%) |  |
| MF | 14 (10%) | 6 (9%) |  |
| MM | 17 (12%) | 3 (5%) |  |
| Others | 12 (8%) | 5 (8%) |  |
| **Donor type, n (%)** |  |  | 0.057<sup>2</sup> |
| MRD 10/10 | 29 (20%) | 20 (30%) |  |
| MUD 10/10 | 90 (62%) | 42 (63%) |  |
| MMUD 9/10 | 27 (18%) | 5 (8%) |  |
| **CMV serostatus, n (%)** |  |  | 0.70<sup>2</sup> |
| R-/D- | 55 (38%) | 20 (30%) |  |
| R-/D+ | 18 (12%) | 8 (12%) |  |
| R+/D- | 22 (15%) | 11 (16%) |  |
| R+/D+ | 51 (35%) | 28 (42%) |  |
| **EBV serostatus, n (%)** |  |  | 0.56<sup>2</sup> |
| R-/D- | 1 (1%) | 2 (3%) |  |
| R-/D+ | 12 (8%) | 3 (5%) |  |
| R+/D- | 14 (10%) | 6 (9%) |  |
| R+/D+ | 111 (76%) | 51 (80%) |  |
| R+/Unknown | 8 (6%) | 2 (3%) |  |
| **ATG received, n (%)** | 146 (100%) | 48 (72%) | <0.001<sup>2</sup> |

**Figure 1.**
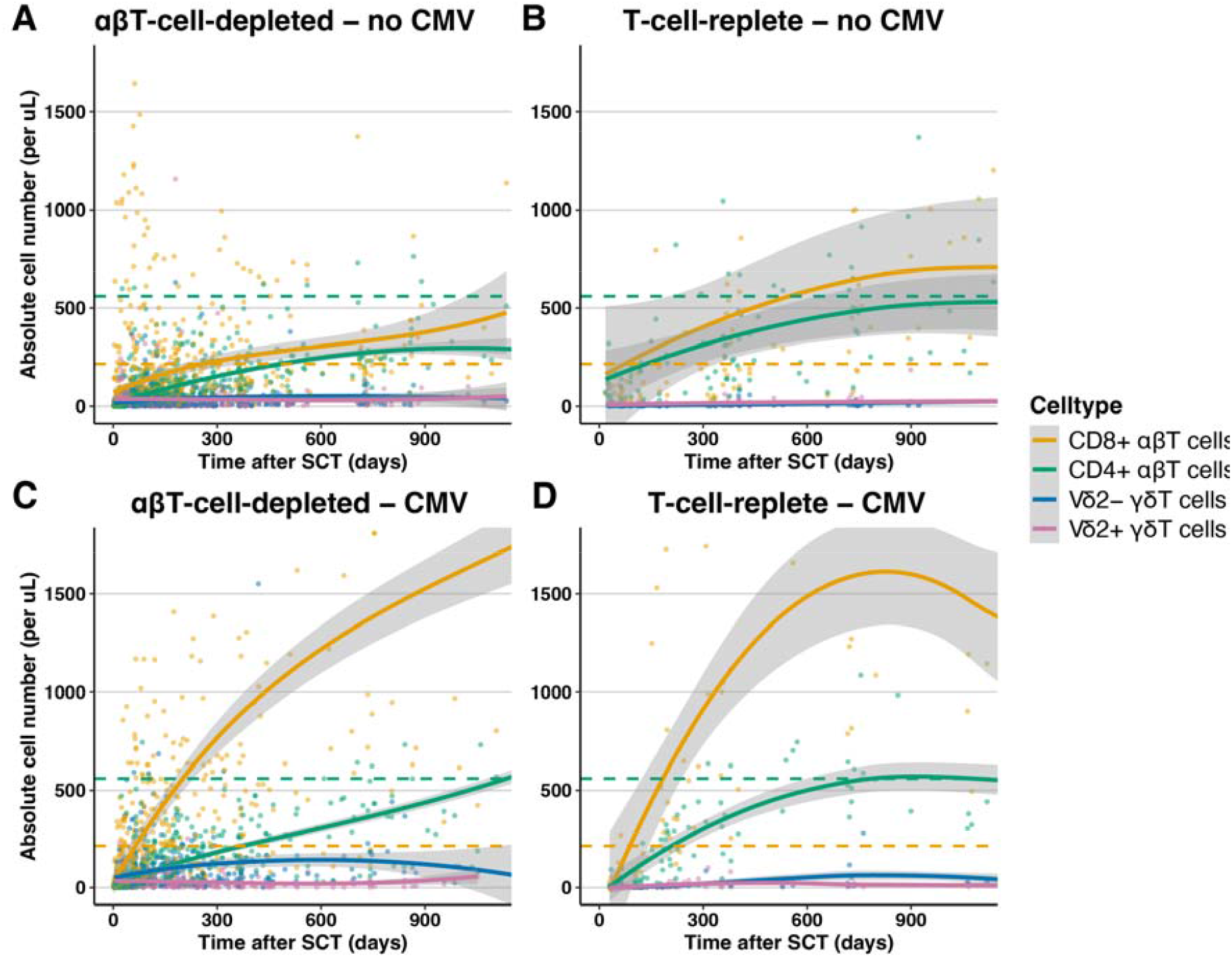
T cell reconstitution after αβT-cell-depleted allo-HSCT and T-cell-replete allo-SCT, stratified by CMV reactivation. Numerical reconstitution of CD8+ αβT cells (ref values 216 – 499), CD4+ αβT cells (ref values 560 – 1067), Vδ2-T cells and Vδ2+ T cells (ref values total γδ T cells 20-120) after αβT-cell-depleted allo-SCT or T-cell replete allo-SCT, stratified by CMV reactivation. **(A)** Patients received αβT-cell-depleted allo-SCT and did not have CMV reactivation (n=83), **(B)** patients received T-cell replete allo-SCT and did not have CMV reactivation (n=36), **(C)** patients received αβT-cell-depleted allo-SCT and had CMV reactivation (n=59), and **(D)** patients received T-cell replete allo-SCT and had CMV reactivation (n=27). All available clinical laboratory data are shown with LOESS curves (span = 1), and 95% confidence intervals are shown. Dashed line: lower limits of reference values for CD8+ αβT cells (orange) and CD4+ αβT cells (green).

### γδ and αβ T cell repertoires follow fundamentally distinct rebuilding rules after αβT cell-depleted transplantation

To define clonal rebuilding dynamics, we performed longitudinal bulk TCR sequencing in a subset of αβT cell-depleted transplant recipients (n = 30). Repertoire evenness was quantified using the D75 index, a repertoire diversity metric defined as the minimum number of unique clonotypes required to account for 75% of the total TCR sequencing reads, with higher values indicating a more diverse and less oligoclonal repertoire. In healthy donors, Vδ2^−^ γδT cells displayed a median D75 of 6.5, whereas Vδ2^+^ γδT cells had a median D75 of 39.5. After transplantation, both γδT subsets exhibited markedly higher early diversity before day 100 (Vδ2^−^ median 60.5; Vδ2^+^ median 90), followed by a significant contraction toward donor-like values beyond day 100 (**Figure 2A-B**; Vδ2^−^ p = 0.008, Vδ2^+^ p = 0.01), indicating an early polyclonal burst followed by repertoire focusing. In contrast, αβT cell repertoires remained persistently restricted. Healthy donors displayed high diversity in CD8^+^ (median D75 221) and CD4^+^ αβT cells (median D75 2394). Post-transplantation, CD8^+^ and CD4^+^ αβT cells exhibited profoundly reduced diversity before day 100 (medians 14.5 and 30.5, respectively), with only minimal diversification thereafter (**Figure 2A-B**). Notably, CD4^+^ αβT cell diversity remained severely constrained even one year post-transplantation. These findings reveal opposing rebuilding strategies: γδT cells undergo early polyclonal expansion followed by clonal focusing, whereas αβT cells, particularly CD4^+^, remain persistently clonally contracted.

**Figure 2.**
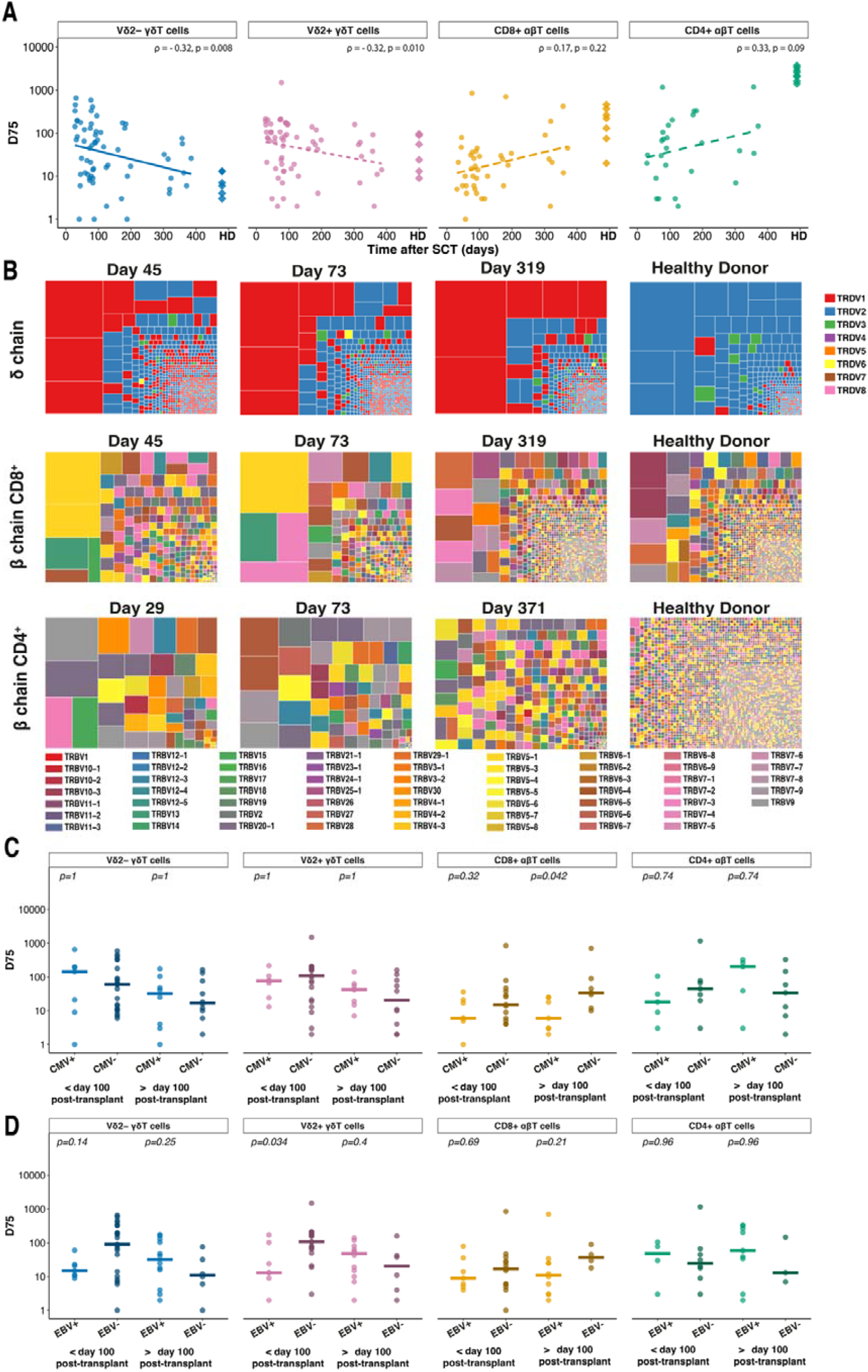
TRD and TRB sequencing were performed to estimate the diversity of the TCR repertoire in αβT-cell-depleted allo-SCT recipients. **(A)** Evenness of TCR repertoire as estimated by unique clonotypes making up 75% of the repertoire (D75). P-values found with Spearman’s correlation, ρ = Spearman’s correlation coefficient. D75 of healthy donors (HD) added for comparison. **(B)** Examples of TCR repertoires of patients vs healthy references (not SCT donors). Delta chain and beta chain CD8^+^ sequential samples from one patient. Beta-chain CD4^+^ representative repertoires from 3 patients. D75/Evenness of TCR repertoire per group (<100 days after SCT, >100 days after SCT) stratified by **(C)** CMV reactivation and **(D)** EBV reactivation. C + D) Median shown. Vδ2^-^ and Vδ2^+^ TRD < 100 days after SCT n = 28 (CMV reactivation n = 7; EBV reactivation n = 7), Vδ2^-^ and Vδ2^+^ TRD > 100 days after SCT n = 18 (CMV reactivation n = 8, EBV reactivation n = 12), CD8^+^ TRB <100 days after SCT n = 20 (CMV reactivation n = 7, EBV reactivation n = 7), CD8^+^ TRB >100 days after SCT n = 14 (CMV reactivation n = 7, EBV reactivation n = 10), CD4^+^ TRB <100 days after SCT n = 14 (CMV reactivation n = 6, EBV reactivation n = 4), CD4^+^ TRB >100 days after SCT n = 13 (CMV reactivation n = 5, EBV reactivation n = 9). P-values found using the Wilcoxon Rank-Sum test, corrected for multiple testing per cell type and group with Holm-Bonferroni correction.

### CMV amplifies γδ and αβ numerically but does not dictate γδ repertoire focusing; EBV associates with early Vδ2^+^ restriction

Despite the marked numerical expansion of Vδ2^−^ γδT cells in CMV-reactivated patients (**Figure 1**), γδTCR repertoire evenness before day 100 did not differ between CMV-reactivated and non-reactivated patients (median D75 143 vs. 60; p = 1; **Figure 2C**). After day 100, Vδ2^−^ γδTCR repertoires became increasingly oligoclonal in both groups (p = 0.008; **Figure 2A**), indicating that repertoire focusing occurred independently of CMV. EBV reactivation was associated with reduced early Vδ2^+^ γδTCR evenness prior to day 100 (median D75 13 vs. 108; p = 0.034 **Figure 2D**), an effect that was not sustained later. This suggests that early Vδ2^+^ repertoire restriction may represent a transient risk state for EBV reactivation. In CD8^+^ αβT cells, CMV reactivation was associated with significantly greater repertoire focusing (median D75 6 vs. 34; p = 0.042; **Figure 2C**), consistent with dominant clonal expansion, whereas CD4^+^ αβT cells were unaffected. EBV had no detectable effect on αβTCR repertoire. Thus, CMV primarily acts as a numerical and clonal amplifier of CD8^+^ αβT cells, while γδTCR repertoire focusing proceeds largely independently of viral reactivation. Additionally, Vδ2^+^ repertoire evenness may modulate susceptibility to antiviral EBV responses.

### αβTCR, but not γδTCR, clonality retains a durable imprint of graft engineering at single-cell resolution

To explore these observations at higher resolution, we performed single-cell RNA and TCR sequencing on donor samples and samples one year after transplantation (EBMT landmark ^47^) in three αβT cell-depleted and three T cell-replete recipients, all of whom experienced CMV reactivation (**Supplementary Figure 2A-B**). After quality control, 95,435 cells were retained, of which 46,223 cells were annotated with a TCR α and/or β chain and 28,716 cells with a TCR γ and/or δ chain (**Supplementary Figure S2C-F**). γδT cell repertoires were not strongly influenced by transplant strategy, as they were highly patient-specific and dominated either by Vδ1^+^ or by Vδ1^−^Vδ2^−^ γδT cell subsets (**Figure 3A, Supplementary Figure S3**). The proportion of single γδT cell clones was low in all patients (<25% of the repertoire). Hyperexpanded γδT cell clones (>75% of the repertoire) were observed in only two patients, one in the αβT cell-depleted (IR067) and one in the T cell-replete (TR007) group (**Figure 3A**). In contrast, the transplant strategy strongly influenced αβTCR-clonality one year after the transplantation. Patients who received αβT cell-depleted transplants showed a markedly higher proportion of hyperexpanded αβT cell clones and a lower proportion of singlet clonotypes compared with T cell-replete recipients (**Figure 3B**). In donor samples, both γδTCR and αβTCR repertoires were comparatively evenly distributed and did not show corresponding skewing (**Supplementary Figure S4A–B**). These data demonstrate that αβT cell clonality, but not γδT cell clonality, retains a durable imprint of graft engineering at single-cell resolution.

**Figure 3.**
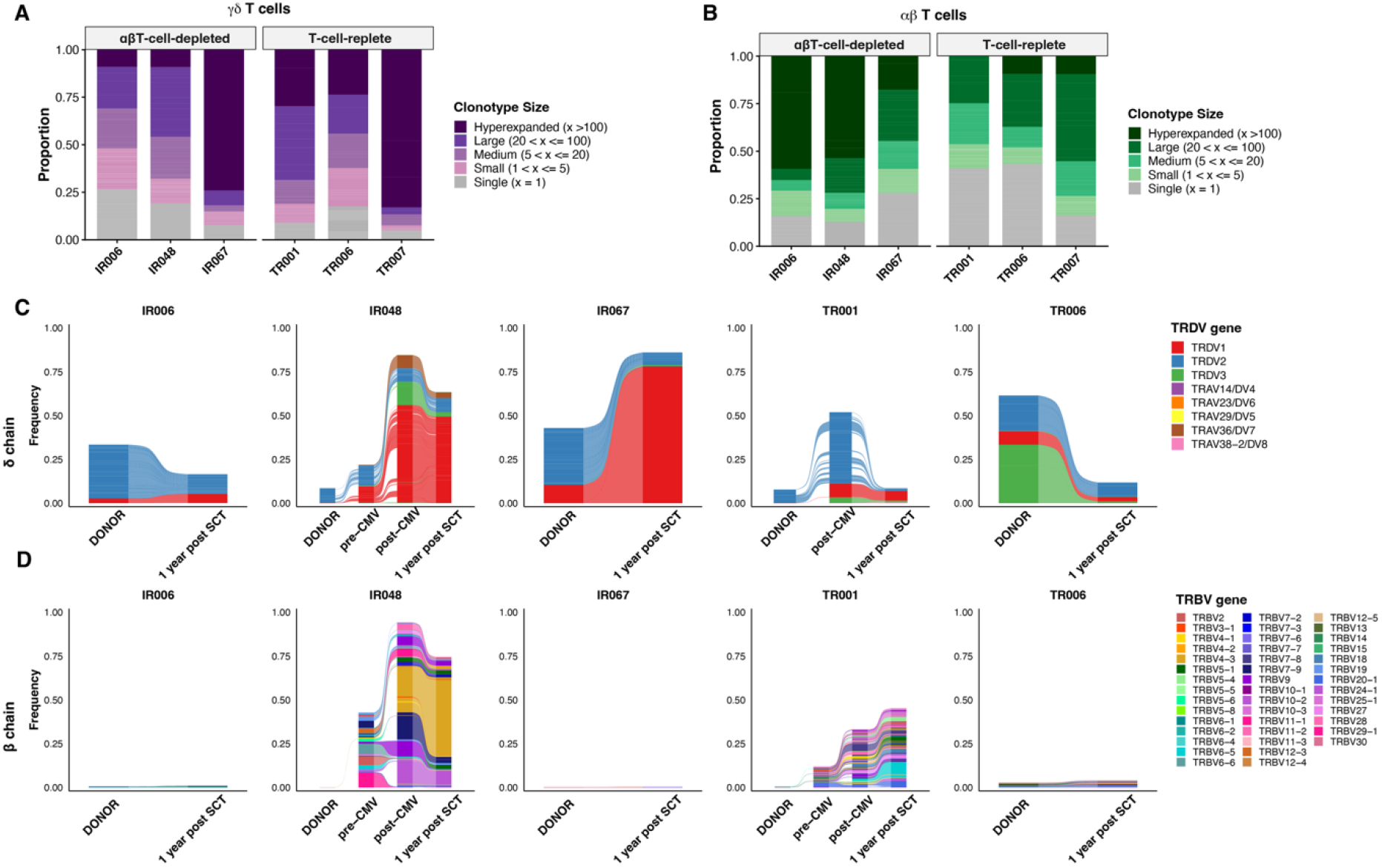
Overview of single-cell TCR data. Single-cell TCR sequencing of peripheral T cells ∼1□year after αβ-T cell–depleted (n□= □3) or T cell–replete (n□= □3) allogeneic SCT. All patients experienced CMV reactivation. Full TCR clones were grouped by size: single (1 cell), small (2–5 cells), medium (6–20 cells), large (21–100 cells) and hyperexpanded (>100 cells). The proportions for each category are shown for individual patients. **A)** γδT cells and **B)** αβT cells. **C-D)** Longitudinal analysis of shared delta chain **(C)** or beta chain **(D)** clonotypes between donors and patients (n = 5), and pre- and post-CMV samples (n=2).

### Vδ2^+^ γδTCRs retain limited donor persistence, whereas Vδ1^+^ γδ and αβ repertoires are largely newly generated

Tracking TCRδ and TCRβ chains between donors and recipients revealed consistent low-level persistence of donor-derived Vδ2^+^ γδTCRs, accounting for <20% of δ chains in recipients one year after transplantation (**Figure 3C**). The majority (>90%) of these shared Vδ2^+^ δ chains were private clonotypes rather than public sequences (**Supplementary Table S2**). In contrast, Vδ1^+^ γδTCR and αβTCR overlap between donors and patients was minimal across platforms, except in rare cases where a small donor clone became hyperexpanded post-transplantation (**Figure 3C**, patient IR067). Overlap of αβTCRs between early post-transplant samples (<100 days) and one-year post-transplant samples was pronounced only after CMV reactivation, with αβTCR overlap exceeding 40% of total β chains in several patients (**Figure 3B**, patients IR048 and TR001). Together, these findings indicate that Vδ2^+^ γδT cells persist partially from the donor pool, whereas the Vδ1^+^ γδ and αβ T cell compartments are predominantly rebuilt de novo after transplantation.

### CMV-driven αβT cell expansion is only partially explained by CMV-specific clonotypes

Considering the substantial increase in CD8^+^ αβT cell numbers in response to CMV reactivation and the expansion of αβT cell clones in αβT cell-depleted allo-HSCT patients, we assessed whether these dominant clones were predominantly CD8^+^ or whether they could also be CD4^+^. In our single-cell RNA sequencing dataset of samples collected 1 year post-allo-HSCT, 8,222 cells expressed either CD8A or CD8B, and 1,669 cells expressed CD4 (**Figure 4A-B**). Because all patients included in this analysis had experienced CMV reactivation, we further examined whether any αβT cell clones were CMV-specific using the Trex package^41,42^. We detected a small proportion of CMV-specific CD8^+^ αβT cell clones in all patients, ranging from 1.8% to 10.5%, with no major differences between αβT cell-depleted and T cell-replete allo-HSCT recipients (Figure 4B). Fewer than 25% of these CMV-specific CD8^+^ clones were single-cell clones, with no differences across transplantation platforms (**Figure 4C**). In contrast, >75% of CMV-specific clones within the CD4^+^ αβT cell compartment were single-cell clones in T cell-replete recipients, whereas CMV-specific CD4^+^ clones were more expanded in patients who received αβT cell-depleted grafts (**Figure 4D**). Interestingly, one patient exhibited a hyperexpanded CMV-specific clone, while the remaining five patients showed similar, lower proportions of CMV-specific clones (4.4–7.2%) (**Figure 4D**). In clones annotated as non-CMV-specific, we observed patterns that closely mirrored those observed in CMV-specific clones. CD8^+^ αβT cell clones were predominantly (hyper)expanded, except in patients IR067 and TR001 (**Figure 4E**), whereas CD4^+^ αβT cell clones were generally smaller and displayed a higher proportion of single-cell clones in the T cell-replete transplant setting (**Figure 4F**). Together, these findings indicate that although CMV reactivation is associated with marked αβT cell expansion, the contribution of CMV-specific clonotypes remains relatively limited one year post-transplantation. Moreover, the proportional contribution of CMV-specific clones is comparable between transplant platforms, despite the CMV-driven αβT cell expansion and the reduction in unique clonotypes observed in αβT cell-depleted transplant recipients.

**Figure 4.**
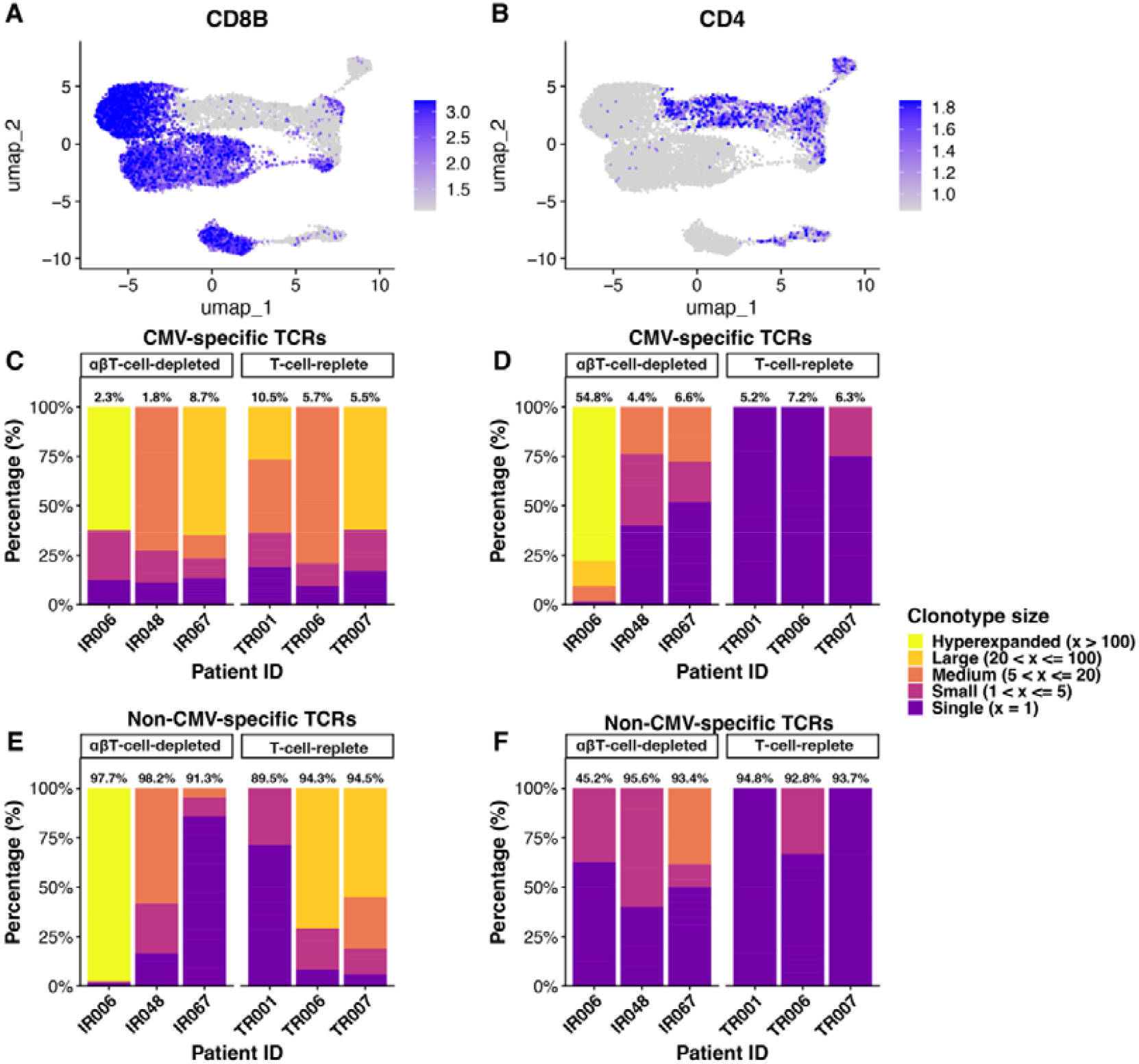
Single cell transcriptomic data and scTCR sequencing data of αβT cells ∼1 year post αβT-cell-depleted allo-SCT (n=3) or T-cell-replete allo-SCT (n=3). **(A)** and **(B)** UMAP including all patient samples showing RNA expression of CD8B (left) and CD4 (right). **(C-F)** Stacked barplot showing **(C)** and **(D)** CMV-specific clones and **(E)** and **(F)** non-CMV-specific clones as determined by the Trex package per patient in CD8+ αβT cells (**C, E**) and CD4+ αβT cells (**D, F**), coloured by TCR clone size. Dark blue = single (1 cell), purple = small (2–5 cells), pink = medium (6–20 cells), orange = large (21–100 cells), and yellow = hyperexpanded (>100 cells).

### Donor and patient T cells occupy distinct transcriptomic states after allo-HSCT

To assess whether patient T cells retained donor-like transcriptional features, we partitioned the scRNA-seq dataset into Vδ2^−^ γδT cells, Vδ2^+^ γδT cells, CD8^+^ αβT cells, and CD4^+^ αβT cells. Pseudo-bulk analysis showed that donors were transcriptionally more similar to one another than to patients across all subsets (**Supplementary Figures S5A–E**). T cellAnnoTator (TCAT) and Azimuth mapping placed post-transplant T cells within a standardized transcriptional reference framework^37,39,40^. Across γδ T cell compartments, patient-derived Vδ2^−^ and Vδ2^+^ γδT cells predominantly exhibited *GZMK*^+^ or *GZMB*^+^ effector phenotypes (**Supplementary Figure S6A–B**). In contrast, donor αβ T cells displayed greater phenotypic diversity, with higher proportions of naïve CD8^+^ and CD4^+^ cells, whereas patient αβ T cells were consistently skewed toward effector memory phenotypes across transplantation settings (**Supplementary Figures S6C–D, S7A–B**).

To further resolve transcriptional structure beyond predefined functional states, we applied non-negative matrix factorization (NMF) at single-cell resolution to define recurrent transcriptional programs across donor and patient T cells (**Supplementary Table S3**). In γδ T cells, cytotoxic effector programs increased after transplantation and CMV reactivation (MP2; **Supplementary Figure S8A**), whereas helper-like programs were highest in donors and reduced after transplantation (MP1; **Supplementary Figure S8A**). In αβ T cells, NMF revealed a shift from donor-enriched naïve programs (MP1; **Supplementary Figure S8B**) toward cytotoxic effector programs (MP2; **Supplementary Figure S8B**). Together, these analyses show that long-term T cell reconstitution after allo-HSCT is characterized by transcriptional states in patients that are distinct from those of their donors across both γδ and αβ lineages.

### Transplantation platform imprints durable gene-expression programs and functional states in T cells

To assess platform-specific effects on T cell transcriptional states, we performed comparative gene-expression analyses between αβT cell-depleted and T cell-replete recipients within the same subsets, identifying four sets of differentially expressed genes (DEGs) (**Supplementary File S1**). No genes were consistently downregulated in the T cell-replete setting (**Figure 5A**); however, seven genes were upregulated across all subsets (*AREG, ELL2, FKBP5, NFKBIA, NR4A2, SELL, and ZFP36*) (**Figure 5B-C**). Increased amphiregulin (*AREG*) expression suggests engagement of tissue-repair or wound-healing programs in T cell-replete recipients^48,49^. Moreover, NMF revealed higher scores of IL7R+ helper signatures in Vδ2^+^ γδT cells of T cell-replete recipients (**Figure 6A**). Together these findings indicate qualitative functional differences between transplantation platforms.

**Figure 5.**
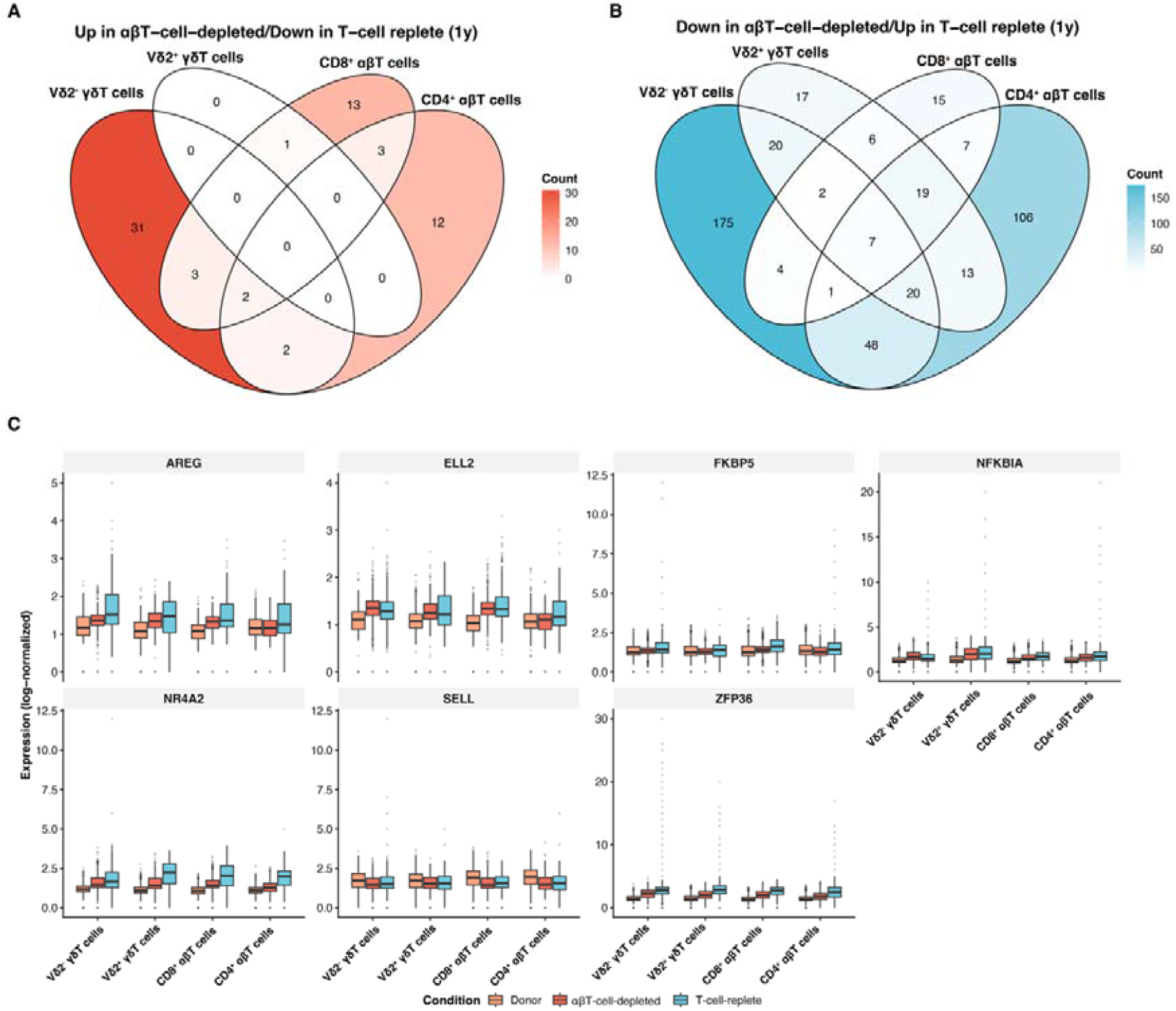
Differential gene expression analysis across T cell subsets at 1 year post-SCT. **(A)** Venn diagram showing the overlap of significantly downregulated genes in T-cell-replete compared to αβT-cell-depleted recipients (log2FC < -1, adjusted p-value < 0.05) across the four T cell subsets **(A)** Venn diagram showing the overlap of significantly upregulated genes in T-cell-replete compared to αβT-cell-depleted recipients (log2FC > 1, adjusted p-value < 0.05) across Vδ2-γδT cells, Vδ2+ γδT cells, CD8+ αβT cells, and CD4+ αβT cells. Numbers indicate the count of differentially expressed genes unique to or shared between cell types. Color intensity reflects the number of overlapping genes.. **(C)** Expression levels of selected differentially expressed genes across cell types and conditions. Boxplots show log-normalized expression values for donor samples (orange), αβT-cell-depleted recipients (red), and T-cell-replete recipients (blue) at 1 year post-SCT.

**Figure 6.**
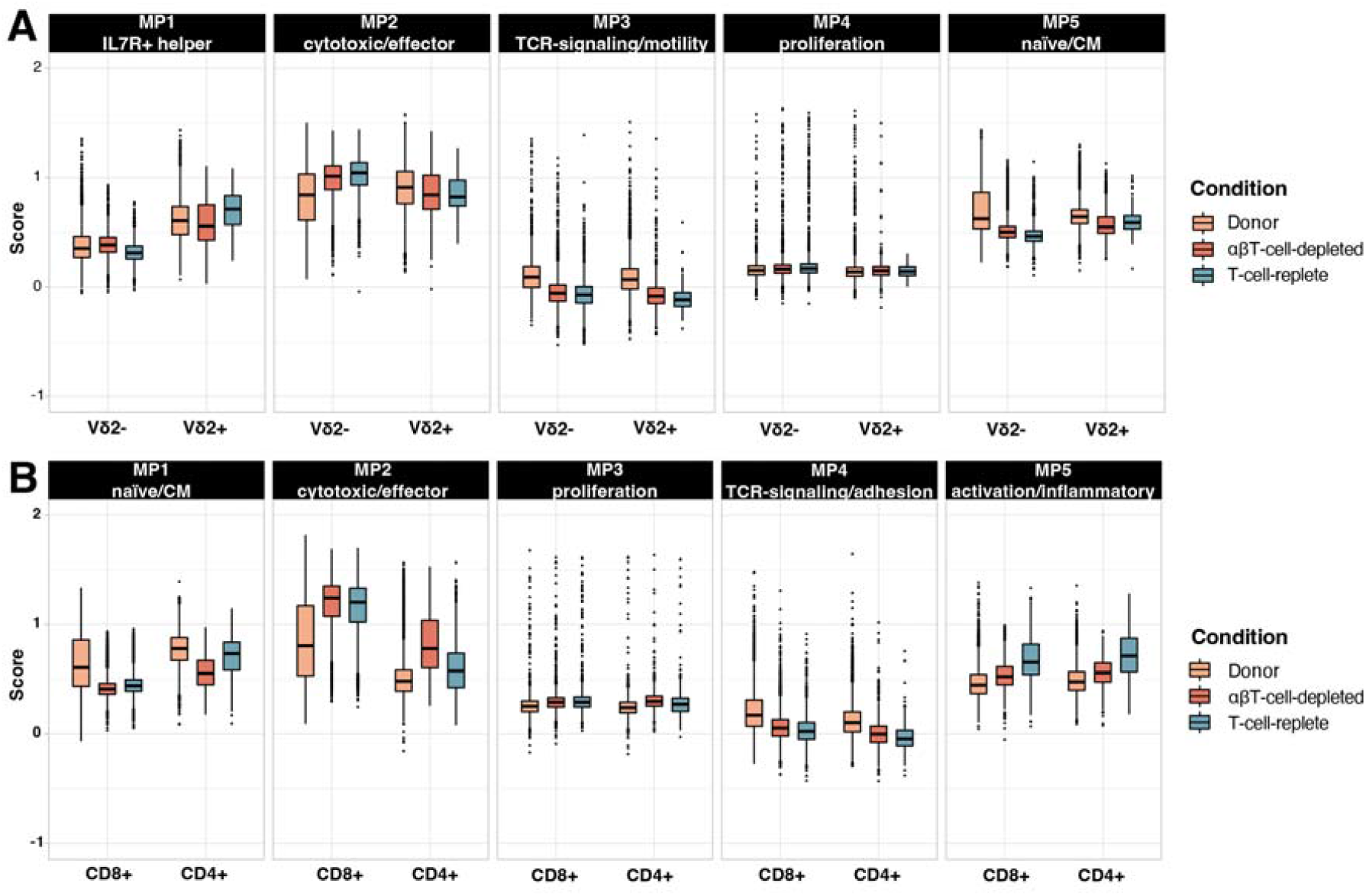
Platform-specific activity of T-cell NMF meta-programs. **(A)** *γδ T cells: Si*ngle-cell module scores of five *γδ T-cell NMF* meta-programs (MP1–MP5) are shown as boxplots, stratified by transplantation platform (donor, *αβT-cell-depl*eted, and T-cell-replete) and *γδT-cell subs*et (V*δ2- and Vδ2+)*. MP1 represents an IL7R^+^ helper-like *γδ T-cell prog*ram (e.g., *IL7R, KLRB1, RORA, IL18RAP*), MP2 a cytotoxic effector program (e.g., *CCL5, NKG7, PRF1, GNLY, FGFBP2*), MP3 a TCR signaling and motility-associated program, MP4 a proliferation/cell-cycle program, and MP5 a naïve/central-memory-like program (e.g., *TCF7, LEF1, CCR7, BACH2*). V*δ2*^−^ and V*δ2*^+^ *γδ T cells are* shown separately. **(B)** αβ T cells: Single-cell module scores of five *αβ T-cell NMF* meta-programs (MP1– MP5) are shown as boxplots, stratified by transplantation platform (donor, *αβT-cell-depl*eted, and T-cell-replete) and αβT-cell subset (CD8+ and CD4+). MP1 corresponds to a naïve/central-memory-like program (e.g., *TCF7, LEF1, IL7R, CCR7*), MP2 to a cytotoxic effector program (e.g., *CCL5, NKG7, PRF1, GNLY*, granzymes), MP3 to a proliferation/cell-cycle program (e.g., *MKI67, PCNA*), MP4 to a TCR signaling and adhesion program (e.g., *FYN, PRKCH, ITGA4, STAT4*), and MP5 to an activation/inflammatory response program (e.g., *TNFAIP3, ZFP36, DUSP1/2, JUN/JUNB, NR4A2, CD69*). CD8^+^ and CD4^+^ *αβ T cells are* shown separately. Boxplots indicate median and interquartile range, with individual points representing single cells.

Functional state scoring using the TCAT pipeline^40^ showed higher cytotoxicity scores in Vδ2^−^ γδT cells from T cell-replete recipients, whereas CD8^+^ and CD4^+^ αβT cells exhibited higher cytotoxicity scores in αβT cell-depleted recipients (**Supplementary Figure S9**), which was confirmed by NMF (MP2; **Figure 6B**). Cytotoxicity scores were consistently higher in patient samples than in donor samples across these subsets (**Figure 6B, Supplementary Figure S10**), indicating that these states are not inherited from donor T cells. Additionally, activation and inflammatory response programs (MP5) were relatively more prominent in αβT cells of T cell-replete recipients (**Figure 6B**).

Collectively, transcriptional analyses demonstrate that the transplantation platform leaves a durable imprint on reconstituting T cells, differentially balancing cytotoxic and tissue-repair–associated programs between γδ and αβT cell lineages.

## Discussion

Allo-HSCT induces a profound and prolonged disruption of immune homeostasis, requiring coordinated rebuilding of both innate-like and adaptive T cell compartments under conditions of lymphopenia, inflammation, and viral exposure. Here, by integrating longitudinal immune phenotyping with bulk and single-cell TCR repertoire and transcriptomic profiling across αβ and γδ T cell subsets, we delineate how graft engineering and viral reactivation jointly shape post-transplant immune architecture over multiple years. By comparing αβT cell–depleted and T cell-replete platforms, we could identify platform-imprinted, lineage-specific patterns of clonal rebuilding, cellular origin, and functional programming that persist long after transplantation.

First, we identified an association between reduced early Vδ2^+^ γδTCR repertoire evenness and subsequent EBV reactivation within the first 100 days post-transplantation. Although correlative, this finding is biologically plausible given the known capacity of Vδ2^+^ γδT cells to recognize EBV-transformed B cells^50^ and genetic variation in BTN- and RhoB-linked regions that affect γδTCR responses^51-54^. In contrast to prior reports linking EBV reactivation to reduced γδT cell numbers^55^, Vδ2^+^ γδT cell counts in our cohort remained stable, suggesting that repertoire composition rather than absolute cell number may be the more sensitive indicator of EBV immune pressure. These data support that early γδTCR repertoire composition could serve as a potential surrogate marker of antiviral immune competence, pending prospective validation.

Second, we demonstrate a significant CMV-driven expansion of Vδ2^−^ γδT cells in αβT cell-depleted transplant recipients, consistent with prior observations in T cell-replete context^10,13,15,18^. Notably, in the αβT cell-depleted platform, this expansion results in a persistent inversion of the Vδ2^+^/Vδ2^−^ ratio and sustained dominance of effector-like Vδ2^−^ γδT cells for up to 3 years post-transplantation. Intriguingly, narrowing of the Vδ2^−^ γδT cell repertoire occurred even in the absence of CMV reactivation, suggesting homeostatic proliferation rather than persistent viral antigen exposure as the primary mechanism underlying repertoire focusing. This aligns with recent findings from healthy CMV-seronegative individuals^19^, and challenges the paradigm of persistent antigen exposure as the sole driver of long-term γδTCR repertoire modulation^18^.

Third, our paired donor–patient single-cell and TCR analyses revealed a strikingly limited overlap of Vδ1^+^ and αβTCR clonotypes between donors and recipients one year after transplantation, independent of transplant platform. This finding indicates that, in contrast to partial donor persistence observed for Vδ2^+^ γδTCRs, the Vδ1^+^ and αβT cell compartment are largely rebuilt de novo after allo-HSCT^56-58^. The minimal donor–patient Vδ1^+^ and αβTCR overlap further supports the notion that post-transplant Vδ1^+^ and αβT cell immunity is not simply a passive extension of the donor repertoire but is actively reshaped by post-transplant antigenic, homeostatic, and inflammatory cues. Importantly, this does not imply equivalent rebuilding dynamics across platforms. Rather, early donor-derived αβT cells present in T cell-replete grafts may condition the post-transplant immune niche through cytokine production, antigen-presenting cell licensing, and competition for homeostatic signals, thereby facilitating more even de novo repertoire diversification despite limited long-term clonotypic persistence^59^.

Fourth, recipients of αβT cell-depleted grafts displayed marked contraction of the αβTCR repertoire with increased hyperexpansion of individual clonotypes. Given that GVHD is a stochastic event, the low incidence observed in this cohort^24^ likely reflects a reduced overall probability of alloreactive encounters rather than a direct relationship with baseline repertoire features. In this context, clonal hyperexpansion may primarily reflect homeostatic filling of available immune space following profound αβT cell depletion, rather than pathogenic alloreactivity. While prior studies in T cell-replete transplantation have linked increased αβTCR diversity and post-transplant clonal hyperexpansion to GVHD risk^60-63^, our data suggest that these relationships are highly platform-dependent. Importantly, the presence of early hyperexpanded clones does not preclude their further expansion in the event of GVHD; rather, it indicates that baseline clonality alone is unlikely to be mechanistically deterministic.

Fifth, despite pronounced CMV-driven expansion of CD8^+^ αβT cells, the fraction of CMV-specific clonotypes one year post-transplantation remained relatively low across platforms, consistent with previous findings^64^. This dissociation between numerical expansion and classical CMV specificity suggests that bystander activation, cytokine-driven proliferation, and NK-like adaptive programs may substantially contribute to CMV-associated CD8^+^ T cell expansion. The preferential expansion of cytotoxic, NK-like CD8^+^ T cells in αβT cell-depleted recipients is consistent with emerging models of CMV-driven adaptive cytotoxic immunity that extend beyond strictly antigen-restricted responses.

Sixth, transcriptomic profiling revealed that both γδ and αβ T cells in patients adopted predominantly effector-like states distinct from those of donors, with the transplantation platform exerting a durable influence on functional polarization. Across T cell subsets, transplantation platform–associated transcriptional programs were characterized by shared upregulation of activation- and stress-responsive genes, whereas they differentially redistributed cytotoxic and tissue-repair– associated states among lineages. In T cell-replete recipients, Vδ2^−^ γδT and αβT cells showed relatively higher engagement of tissue-repair–associated programs, including AREG expression^48,49^, whereas αβT cell-depleted recipients showed preferential skewing of CD8^+^ αβT cells toward cytotoxic effector programs. These divergent transcriptional profiles suggest that graft engineering qualitatively reprograms effector function, potentially balancing tissue protection and cytotoxic immunity in a platform-dependent manner. Whether these programs differentially influence viral clearance, GVHD, or epithelial recovery remains to be determined.

Collectively, our data support a model in which post-transplant immune rebuilding is shaped by three interacting forces: (i) graft engineering, which imposes durable constraints on αβTCR diversity and platform-specific effector programming; (ii) viral exposure, particularly CMV, which acts as a dominant numerical amplifier of cytotoxic lineages without fully dictating long-term repertoire architecture; and (iii) intrinsic homeostatic mechanisms, which drive early polyclonal γδ expansion followed by progressive repertoire focusing. Crucially, these forces act on T cell lineages with fundamentally different relationships to the detectable donor repertoire. Within this framework, Vδ2^+^ γδ T cells constitute a rapidly adaptive compartment that partially persists from the donor pool, whereas Vδ2^−^ γδ and αβT cells undergo substantial post-transplant repertoire reconstitution, characterized by the emergence of new dominant clonotypes not detectable in the graft.

These findings suggest that early post-transplant TCR repertoire composition may represent a previously underappreciated determinant of immune behavior after allo-HSCT. As post-transplant interventions such as DLIs, kinase inhibitors, and immunomodulatory therapies become increasingly integrated into standard care^65,66^, baseline clonal diversity and lineage balance may critically condition both therapeutic efficacy and immune toxicity. Likewise, early restriction of γδT cell repertoires may influence antiviral control and responsiveness to emerging γδ-directed strategies^14^. These data support the incorporation of TCR repertoire monitoring into post-transplant risk-stratification frameworks^66^. Moreover, genetic variation in γδTCR ligands, such as BTN2A1, BTN3A1, and RhoB, may further condition Vδ2^+^ γδT cell responsiveness ^54^.

Several limitations warrant consideration. The number of patients analyzed by single-cell RNA and TCR sequencing was small, limiting statistical power for transcriptomic and clonotype-level comparisons and precluding robust adjustment for conditioning, disease type, and GVHD prophylaxis. In addition, matched bulk and single-cell TCR sequencing was not available for all patients. Where overlap existed, concordant clonotype detection across platforms supports the robustness of the analytical approach. Larger, prospectively designed studies will be required to validate the clinical implications of these findings and to define their relevance for viral control, GVHD, and relapse across additional transplantation strategies^67^.

In conclusion, post-transplant immune reconstruction is not uniform but is durably shaped by graft engineering, viral exposure, and intrinsic homeostatic forces in a lineage-specific manner. By defining distinct rebuilding rules for γδ and αβT cells at numerical, clonal, and functional levels, these findings provide a framework for understanding individual immune trajectories after allo-HSCT. As immune-directed therapies continue to expand, such insights may enable more precise alignment of post-transplant interventions with each patient’s evolving immune architecture.

## Acknowledgments

The authors would like to thank the Single Cell Genomics Core Facility at the Princess Máxima Center for Pediatric Oncology for technical support.

## Author contributions

Conceptualization: MdW, JK; Methodology: AS, PB, AJ, MdW, JK; Formal analysis: AS, PB, AJ; Unique resources: JY, HTS, HS, SP; Data collection: AJ, EK, DdB, MN, LG; Investigation: all authors; Writing; original draft: AS, PB, MdW, JK; Writing; editing and revision: all authors; Supervision: ZS, MdW, JK.

