## Supplementary Methods and Figures for "Platform-Imprinted Transcriptional and Clonal Remodeling of αβ and γδT Cells After Allogeneic Transplantation"

**Details of the Clinical Cohort**

Patients with hematological malignancies who underwent αβ T-cell-depleted stem cell transplantation at the University Medical Center Utrecht between 2015 and 2017 were included in this retrospective analysis. The conditioning regimen consisted of ATG (Thymoglobulin®) 1.5 mg/kg i.v. on days −12 to −9; fludarabine 40 mg/m² i.v. on days −5 to −2; and busulfan i.v. (Busilvex®) with AUC90 on days −5 to −2. αβ T cell reduction was performed by depletion using anti-αβ T cell receptor (TCR) antibodies in combination with magnetic microbeads on the automated CliniMACS device (Miltenyi Biotec, Bergisch Gladbach, Germany), as described previously [29] and according to standard operating procedures. The maximum allowable residual αβ T-cell number was 5 × 10⁵ cells/kg. The allograft was infused within 24 hours after depletion. Mycophenolate mofetil (MMF) was administered at 15 mg/kg three times daily (maximum 3000 mg total) for 28 days as single GVHD prophylaxis. Patients received a pre-emptive donor lymphocyte infusion (DLI; 1 × 10⁵ CD3⁺ T cells/kg) at three months after stem cell infusion, in accordance with guidelines for T cell depletion, provided patients were free of GVHD and not receiving immunosuppressive medication at the time of DLI.

As a comparator group, patients who underwent a T-cell-replete stem cell transplantation at UMCU between 2011 and 2018 were included. Patients received either reduced-intensity conditioning (RIC) or myeloablative conditioning (MA). RIC regimens consisted of: (i) fludarabine 30 mg/m² i.v. on days −3 to −1 with total body irradiation (TBI) 2 Gy on day 0, with or without ATG (Thymoglobulin®) 2 mg/kg i.v. on days −7 to −4; (ii) fludarabine 30 mg/m² i.v. on days −6 to −2 with melphalan 70 mg/m² i.v. on days −3 to −2, with or without ATG 2 mg/kg i.v. on days −9 to −7; or (iii) fludarabine orally 50 mg/m² on days −9 to −4 with melphalan 100 mg/m² i.v. on day −3 and alemtuzumab 15 mg on days −3 and −2. MA regimens consisted of either cyclophosphamide 60 mg/kg i.v. on days −4 to −3 with TBI 6 Gy on days −1 to 0, with or without ATG 2 mg/kg i.v. on days −8 to −5; or fludarabine 40 mg/m² i.v. on days −5 to −2 with busulfan (Busilvex®) i.v. on days −5 to −2 and ATG 1.5 mg/kg i.v. on days −9 to −6. In most regimens, MMF was administered at 15 mg/kg three times daily (maximum 3000 mg total) for 84 days, in combination with cyclosporine (2 × 1.5 mg/kg i.v. or 4.5 mg/kg p.o.) for 180 days as GVHD prophylaxis. In melphalan-containing regimens, MMF was used as single GVHD prophylaxis at the same dosage. In alemtuzumab-containing regimens, cyclosporine was used as single GVHD prophylaxis. Peripheral blood stem cells were used as the graft source in both cohorts. At regular time points after allo-HSCT, levels of B cells, T cells, and NK cells, as well as their subsets, were measured as part of routine clinical care. CMV reactivation was defined as a viral load >250 IU/mL and EBV reactivation as >1000 IU/mL, as assessed by PCR. Data regarding immune reconstitution and viral reactivations were retrieved from the hospital medical record system.

**Next-generation sequencing of the T cell receptor**

Frozen PBMC samples from patients and healthy donors were thawed. PBMCs were stained with the following monoclonal antibodies: anti-CD3 eFluor450 (clone OKT3, eBioscience), anti-TCRαβ APC (clone IP26, eBioscience), anti-TCRγδ PE (clone IMMU510, Beckmann Coulter), anti-CD8 PerCP-Cy5.5 (clone RPA-T8, Biolegend), and anti-CD4 FITC (clone RPA-T4, eBioscience). Next, cells were sorted (BD FACSAriaTM II cell sorter) into CD8^+^ and CD4^+^ abT cell fractions (CD3pos, TCRαβpos, CD8pos or CD4pos) and gdT cell fractions (CD3pos, TCR γδpos). After sorting, total RNA was isolated using RNeasy Mini or RNeasy Micro kits (QIAGEN) according to the manufacturer’s instructions, with kit selection based on input cell number (RNeasy Mini for >5×10^5 cells; RNeasy Micro for ≤5×10^5 cells).

The protocol for sequencing the variable region of the β chain of the T cell receptor (TCR) for the CD4 and CD8 fractions was adapted from Mamedov *et al.* [1]. cDNA was synthesized (SuperScriptTM II Reverse Transcriptase, Thermo Fisher Scientific) utilizing a specific primer at the 3’ constant region and a universal template switch adaptor at the 5’ end of the V region (Supplementary Table S1). Purified cDNA (NucleoSpin Gel and PCR Clean-UP, Macherey-Nagel) was amplified using Q5® High-Fidelity DNA polymerase (New England BioLabs, Inc.) on a T100TM Thermal Cycler (Bio-Rad). A specific nested primer located in the constant region and a step-out primer, which anneals to the switch adaptor, were used (Supplementary Table S1). The PCR product was loaded onto a 1.5% agarose gel, electrophoresed, and products between 400-600 base pairs were size-selected and purified (NucleoSpin Gel and PCR Clean-UP, Macherey-Nagel).

A multiplex PCR was used to sequence the variable region of the δ-chain TCR for the γδ fraction. First cDNA was synthesized (SuperScriptTM II Reverse Transcriptase, Thermo Fisher Scientific) and purified (NucleoSpin Gel and PCR Clean-UP, Macherey-Nagel). cDNA was amplified using using Q5® High-Fidelity DNA polymerase (New England BioLabs, Inc.) on a T100TM Thermal Cycler (Bio-Rad) and purified (NucleoSpin Gel and PCR Clean-UP, Macherey-Nagel).

Both the PCR products of the β and the δ chain were loaded on the QIAxcel (Qiagen) and the intensity of the PCR product was measured as relative fluorescent units (RFU). Based on the RFU the samples were diluted for the HTS library preparations, which were carried out using the HTSgo-LibrX kit with HTSgo-IndX indices following the manufacturer’s (Gendx) recommendations. Subsequently, samples were purified using HighPrep PCR beads (GC Biotech), and HTS was performed on an Illumina MiSeq 500 (2 x 250 bp read length).

**NGS data analysis**

Sequences of the CDR3 region of the β and the δ chain of the TCR were extracted from the raw Illumina files using MiXCR software[2]. Only productive reads were considered, and sequences with a read count <2 were excluded from analysis. To correct for potential bias, read counts were normalized to the sample with the lowest count (8400 reads). Repertoire analyses were performed using the immunarch package (version 0.9.0) in R version 4.2.2 [3].

**Single-cell RNA sequencing and single-cell TCR sequencing**

Frozen PBMC samples from patients collected ~1 year after SCT (range 356–463 days) were thawed. All patients had prior CMV reactivation. In addition, corresponding donor PBMC samples were available for 5 patients, and for 2 of these patients, we included paired pre- and post-CMV reactivation samples. PBMCs were stained with the following monoclonal antibodies: anti-CD3 Pacific Blue (clone UCHT1, BD), anti-TCRαβ APC (clone IP26, eBioscience), anti-TCRγδ PE-Cy7 (clone IMMU510, Beckmann Coulter), anti-Vδ1 PE (clone REA173, Miltenyi), anti-Vδ2 FITC (clone B6, Biolegend), and anti-CD8 PerCP-Cy5.5 (clone RPA-T8, Biolegend). Subsequently, they were stained with LIVE/DEAD Aqua (ThermoFisher). Next, cells were sorted (BD FACSAriaTM II cell sorter) into abT cell fractions (alive cells, CD3pos, TCRαβpos) and gdT cell fractions (alive cells, CD3pos, TCR γδpos).

Libraries for single-cell transcriptome sequencing and scTCR-seq were prepared using the Chromium Single-Cell 5′ Library GEM-X Gel Bead and Construction Kit and Chromium Single-Cell V(D)J Enrichment Kit (v3, 10x Genomics, CA, USA). Custom primers for the capture of γδTCR transcripts were produced as described by Tan et al [4]. The scRNA-seq and scTCR-seq libraries were sequenced on the Illumina Novaseq 6000 platform.

**scRNAseq expression data analysis**

Raw FASTQ files from 5′ gene expression (GEX) and targeted V(D)J libraries were merged by lane for each sample and organized into multimodal inputs. Joint processing of matched GEX and V(D)J data was performed with Cell Ranger (v8.0.1) using *cellranger multi* with GRCh38 references for gene expression and V(D)J. To assign cells to their donor of origin, SNPs were called from the Cell Ranger BAM using *cellSNP-*lite [5], with a gnomAD hg38 variant panel, and genotype-based demultiplexing was performed with *Vireo* [6], assuming two donors per library; singlet, doublet, and unassigned calls were added to the Seurat metadata. Seurat (5.1.0) was used in R version 4.4.1 for dimensional reduction, cell clustering, and differential gene expression analysis [7-10]. Azimuth was used to integrate our dataset with CD8+ abT cell, CD4+ abT cell, and gdT cell subsets from the Human Immune Health Atlas by the Allen Institute [9, 11]. Furthermore, we used the T-CellAnnoTator (TCAT) pipeline to characterize functional programs in T-cells [12].

**scTCRseq data analysis**

Alignment of gdTCRs was performed by using IMGT/HighV-QUEST on unfiltered FASTA outputs from CellRanger [13]. Unproductive sequences and duplicated barcodes were filtered out. scTCRseq data was added as metadata to the scRNAseq Seurat objects by matching the barcodes. We defined single clones as one cell with a unique TCR, small clones as one to five cells with the same TCR, medium clones as five to twenty cells with the same TCR, large clones as twenty to one hundred cells with the same TCR, and hyperexpanded clones as over one hundred cells with the same TCR. abTCRs were screened for CMV specificity using the scRepertoire and Trex packages [14, 15], which annotate αβ TCRs with epitope data from VDJdb, McPAS-TCR, IEDB, and PIRD databases [16-19].

**Non-negative matrix factorization (NMF) analysis**

NMF was performed as described earlier by Gavish et al. [20], independently for each sample to capture within-sample transcriptional heterogeneity while minimizing sensitivity to batch effects. Factorization was run across multiple ranks (k = 4–9), and for each resulting program, the top 50 genes ranked by factor loadings were retained.

To derive robust and non-redundant programs, a multi-step filtering strategy was applied. First, within-sample robustness was achieved by retaining only programs that recurred across multiple k values within the same sample, defined as≥70% overlap in their top genes. Second, across-sample robustness was imposed by requiring programs to share ≥20% of their genes with at least one program identified in another sample. Third, to remove redundancy within samples, overlapping programs were ranked by their maximal cross-sample overlap, and only the highest-ranking program was retained when overlaps exceeded 20%.

Robust programs were subsequently grouped into higher-order meta-programs using a greedy gene-overlap–based clustering approach, as previously described. Programs were iteratively clustered based on Jaccard similarity of their gene sets, and each meta-program (MP) was represented by a consensus gene signature reflecting the most frequently shared genes across its constituent programs. For both αβ and γδ T cells, this approach yielded five conserved MPs. MP activity was quantified by computing module scores at the single-cell level (via the *AddModuleScore* function of Seurat), which were then compared across donors and transplantation platforms.

**Statistical analyses**

Statistical analyses were performed using R (R version 4.2.2). Wilcoxon rank-sum test was used to test for differences between groups as described in figure legends. Statistical analyses were performed in R (version 4.2.0) using the nlme package. For the αβT‑cell–depleted cohort, we fitted a piecewise linear mixed‑effects model with a single knot at 60 days post‑transplant. The model included fixed effects for time before the knot (Days1), time after the knot (Days2), CMV status, and their two interaction terms (Days1×CMV and Days2×CMV). Patient ID was entered as a grouping factor supplying random intercepts and random slopes to account for within‑patient correlations over time. For the T‑cell–replete cohort, we fitted a simpler linear mixed‑effects model with a single slope: fixed effects were time (continuous), CMV status, and their interaction, and patient ID entered as a random intercept. We used two‑sided Wald t‑tests on the time × CMV interaction to evaluate effect of CMV reactivation on reconstitution rates. P-values < 0.05 were considered significant.

**Data availability**

The single-cell RNA-seq data generated in this study are not publicly available due to patient privacy restrictions, but can be accessed upon reasonable request from the corresponding author.

**Supplementary Tables**

**Supplementary Table S1. Impact of CMV reactivation on the ratio of Vδ2^+^/Vδ2^-^ γδT cells.** N indicates the total number of samples.

| Platform | CMV | Ratio < day 100 (IQR) | n | Ratio > day 100 (IQR) | n |
| --- | --- | --- | --- | --- | --- |
| **αβT-cell-depleted** | No | 1.80 (0.66-4.18) | 239 | 0.61 (0.22-1.42) | 260 |
| **αβT-cell-depleted** | Yes | 0.58 (0.25-1.66) | 180 | 0.16 (0.07-0.39) | 158 |
| **T-cell-replete** | No | 5.62 (5.11-9.58) | 15 | 1.63 (0.28-5.62) | 52 |
| **T-cell-replete** | Yes | 1.49 (0.91-11.33) | 7 | 0.40 (0.20-2.20) | 47 |

**Supplementary Table S2. Overview of private clones shared between donor and allo-HSCT recipient**

| **PatientID** | **Private shared Vδ2+ δ chains (%)** |
| --- | --- |
| IR006 | 94,2 |
| IR048 | 99,5 |
| IR067 | 96,3 |
| TR001 | 96,8 |
| TR006 | 100 |

Definition private: not present in database with public clones (Vyborova et al., 2022)

**Supplementary Figures**

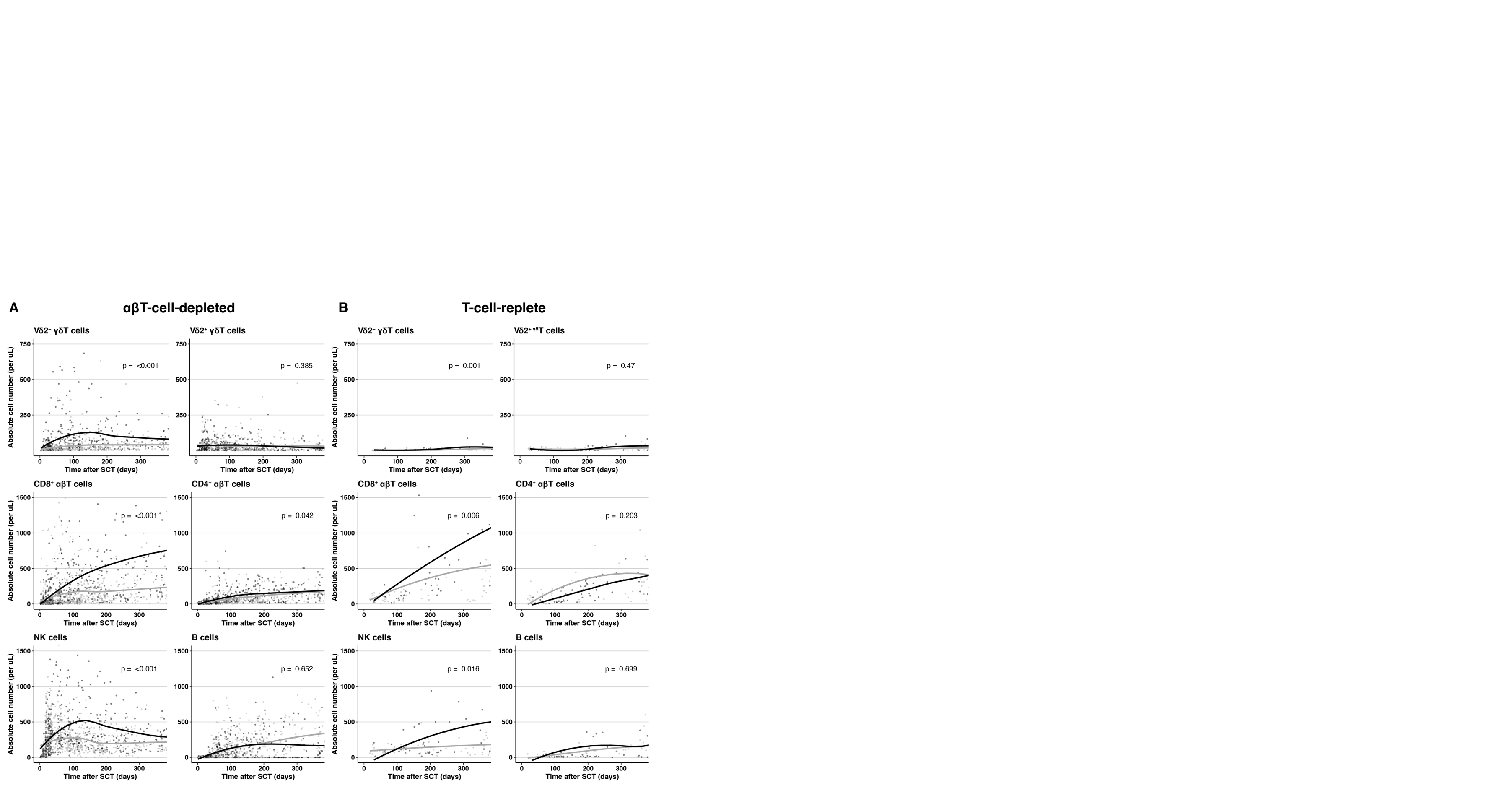

***Supplementary Figure S1. Impact of CMV reactivation on immune reconstitution. (A)*** *Immune reconstitution of Vδ2^-^ γδT cells, Vδ2^+^ γδT cells, CD8^+^ αβT cells, CD4^+^ αβT cells, NK cells, and B cells after αβT-cell-depleted allo-SCT, stratified by CMV reactivation. Black = CMV reactivation (n=59), grey = no CMV reactivation (n=83). P-values show whether CMV had a significant impact on the slope in the first 60 days post-transplantation using a linear mixed-effects model. LOESS curves shown, span = 0.9.* ***(B)*** *Immune reconstitution of Vδ2^-^ γδT cells, Vδ2^+^ γδT cells, CD8^+^ αβT cells, CD4^+^ αβT cells, NK cells, and B cells after T-cell-replete allo-SCT, stratified by CMV reactivation. Black = CMV reactivation (n=27), grey = no CMV reactivation (n=36). P-values show whether CMV had a significant impact on the slope using a linear mixed-effects model. LOESS curves shown, span = 0.9.*

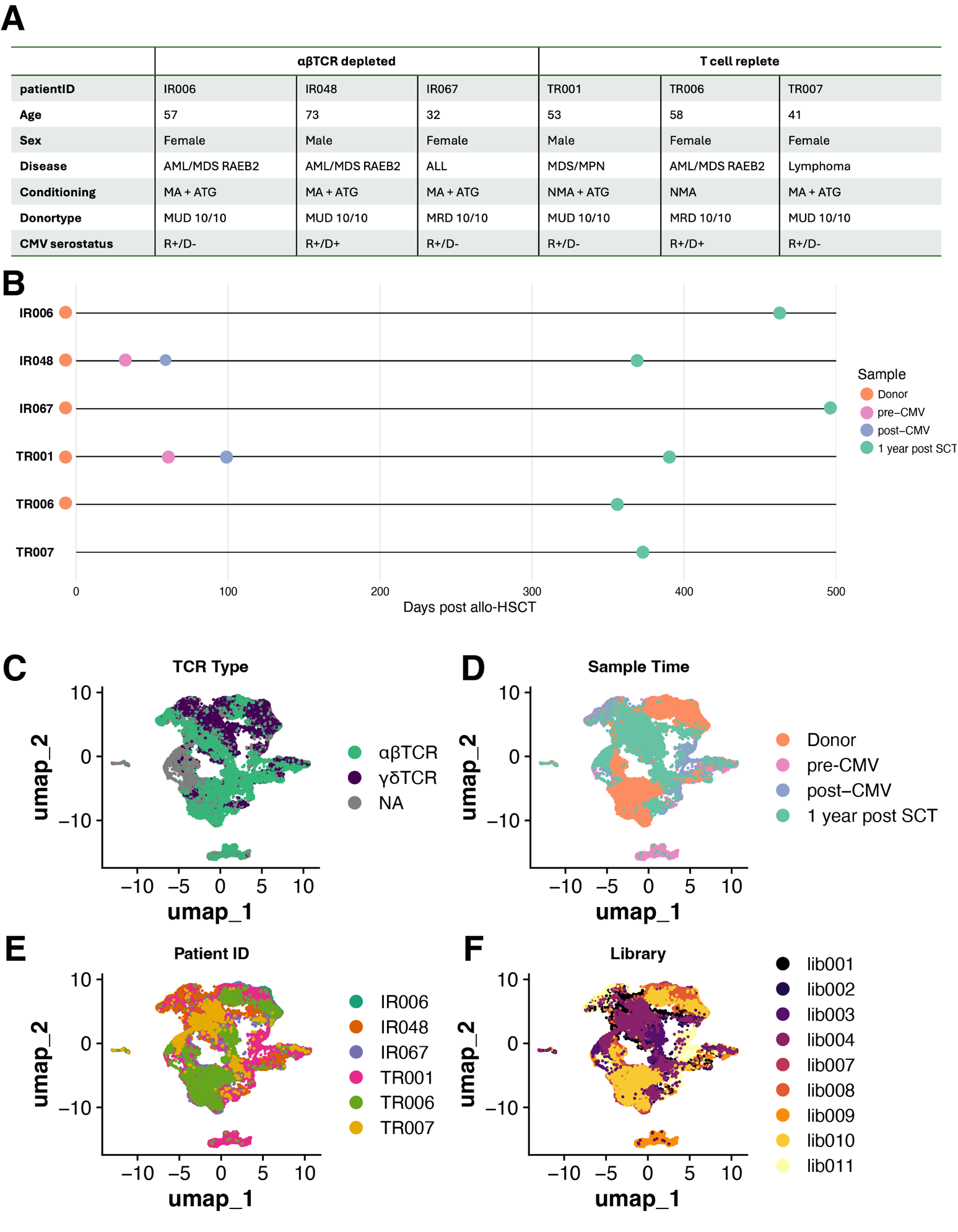

***Supplementary Figure S2. Overview of patient samples used for scRNAseq and scRNAseq data. (A)*** *Overview of patient characteristics included for scRNA sequencing. AML = Acute Myeloid Leukemia, MDS RAEB-2 = Myelodysplastic Syndrome, Refractory Anemia with Excess Blasts type 2, ALL = Acute Lymphoblastic Leukemia, MA = myeloablative, NMA = non-myeloablative, ATG = anti-thymocyte globulin, MUD = matched unrelated donor, MRD = matched related donor, CMV = cytomegalovirus, R = recipient, D = donor.* ***(B)*** *Timeline of samples.* ***(C-F****) UMAPs of sequenced cells, colored by* ***C)*** *TCR,* ***D)*** *time of sampling,* ***E)*** *Patient ID, and* ***F)*** *Sequencing Library. Cells were clustered after filtering out V(D)J genes. A total of 95435 cells were retained after QC, of which 46223 were annotated with a TCR α and/or β chain, and 28716 cells were annotated with a TCR γ and/or δ chain. Cells annotated with more than one chain type were classified as doublets and excluded from downstream analysis. No batch correction was performed****.***

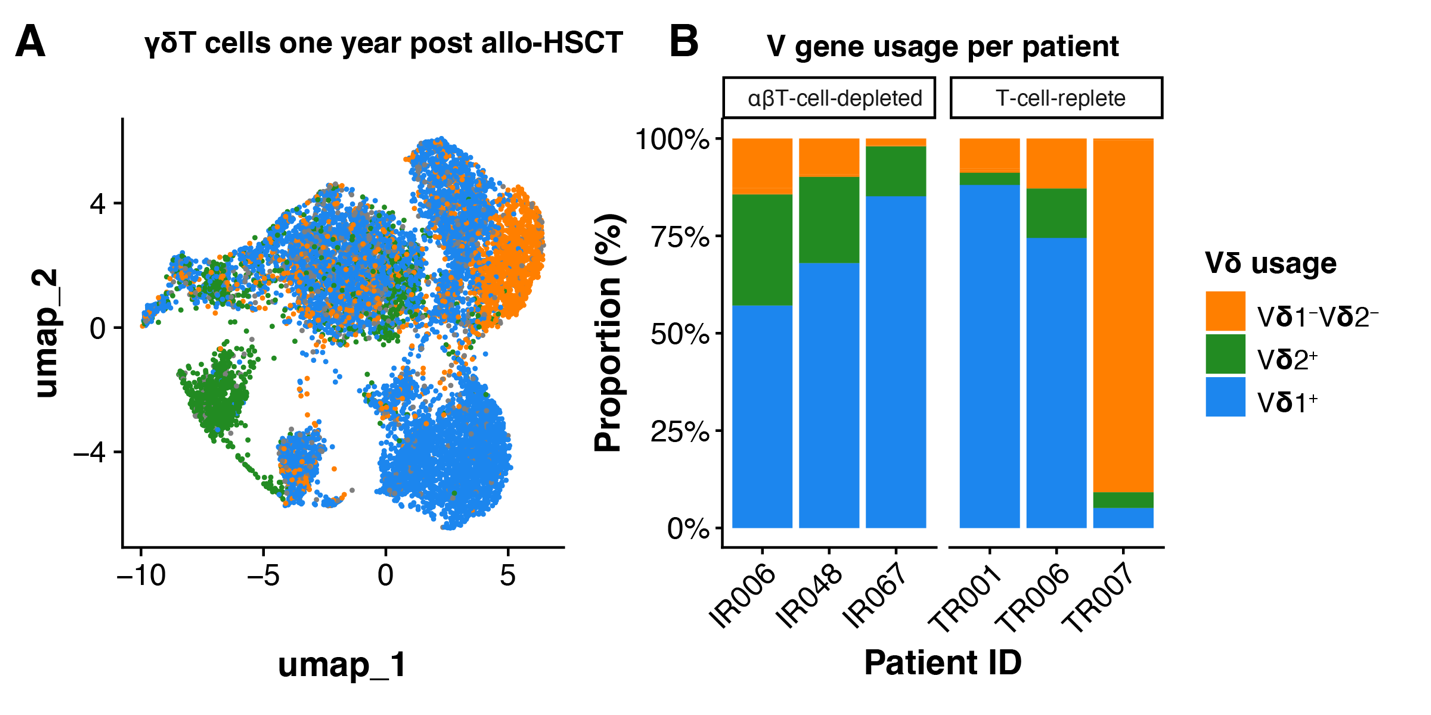

***Supplementary Figure S3. Single-cell TCR sequencing analysis of V-gene usage in γδ T cells 1 year after allo-SCT****. All patients experienced CMV reactivation* ***(A)*** *UMAP of annotated γδ T cells from all patient samples, colored by V gene usage: Vδ1⁺ (blue), Vδ2⁺ (green), and Vδ1⁻Vδ2⁻ (orange). No batch correction was performed.* ***(B)*** *Stacked bar plots showing the proportion of each Vδ subset per patient.*

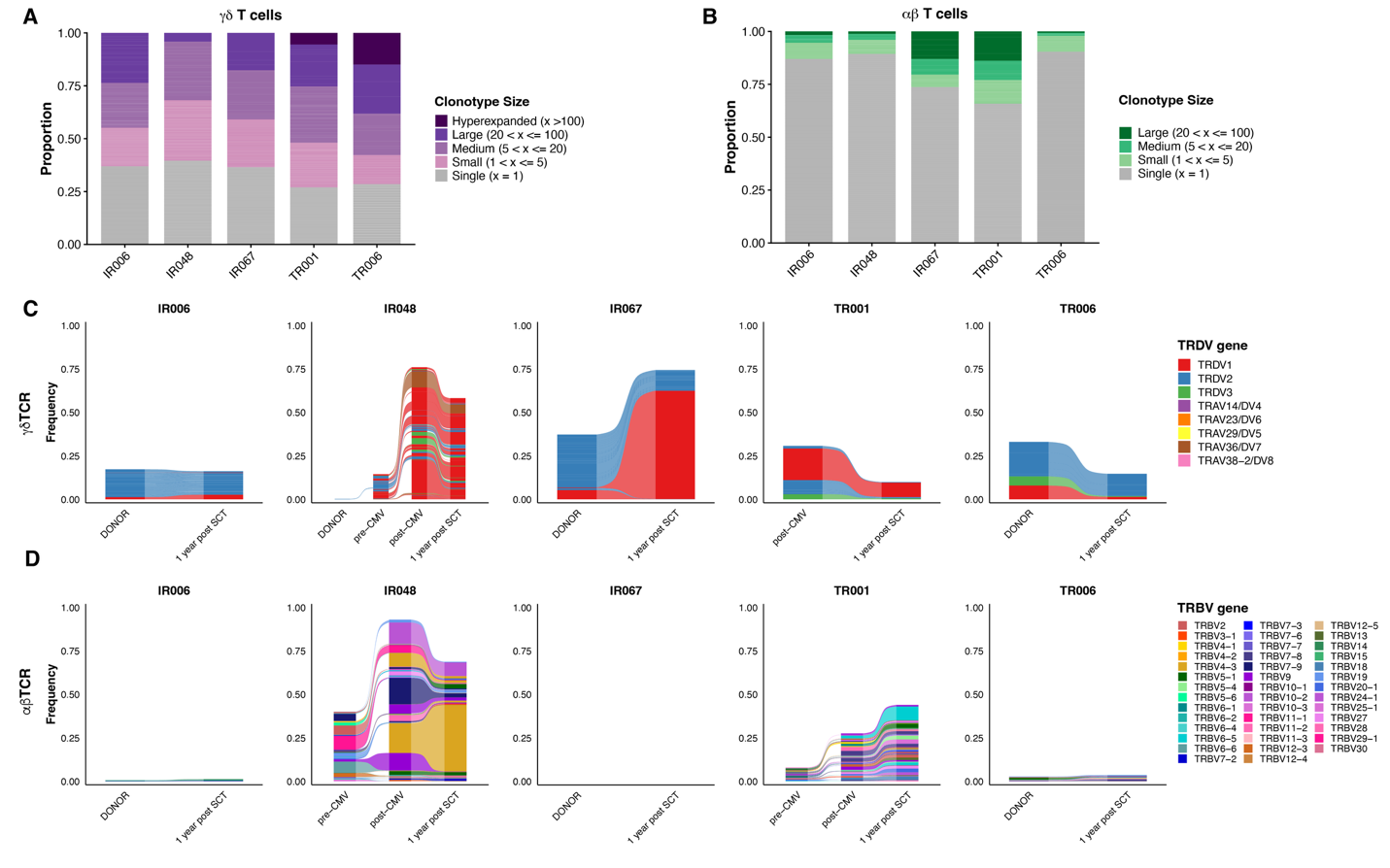

***Supplementary Figure S4. Overview of single-cell TCR data A-B)*** *Single‐cell TCR sequencing of peripheral T cells of stem cell transplantation donors (n=5). Full TCR clones were grouped by size: single (1 cell), small (2–5 cells), medium (6–20 cells), large (21–100 cells) and hyperexpanded (>100 cells). The proportion of each category is shown for individual donors.* ***(A)*** *γδT cells and* ***(B)*** *αβT cells.* ***(C-D)*** *Longitudinal analysis of shared γδTCR* ***(C)*** *or αβTCR* ***(D)*** *clonotypes between donors and patients (n = 5), and pre- and post-CMV samples (n=2).*

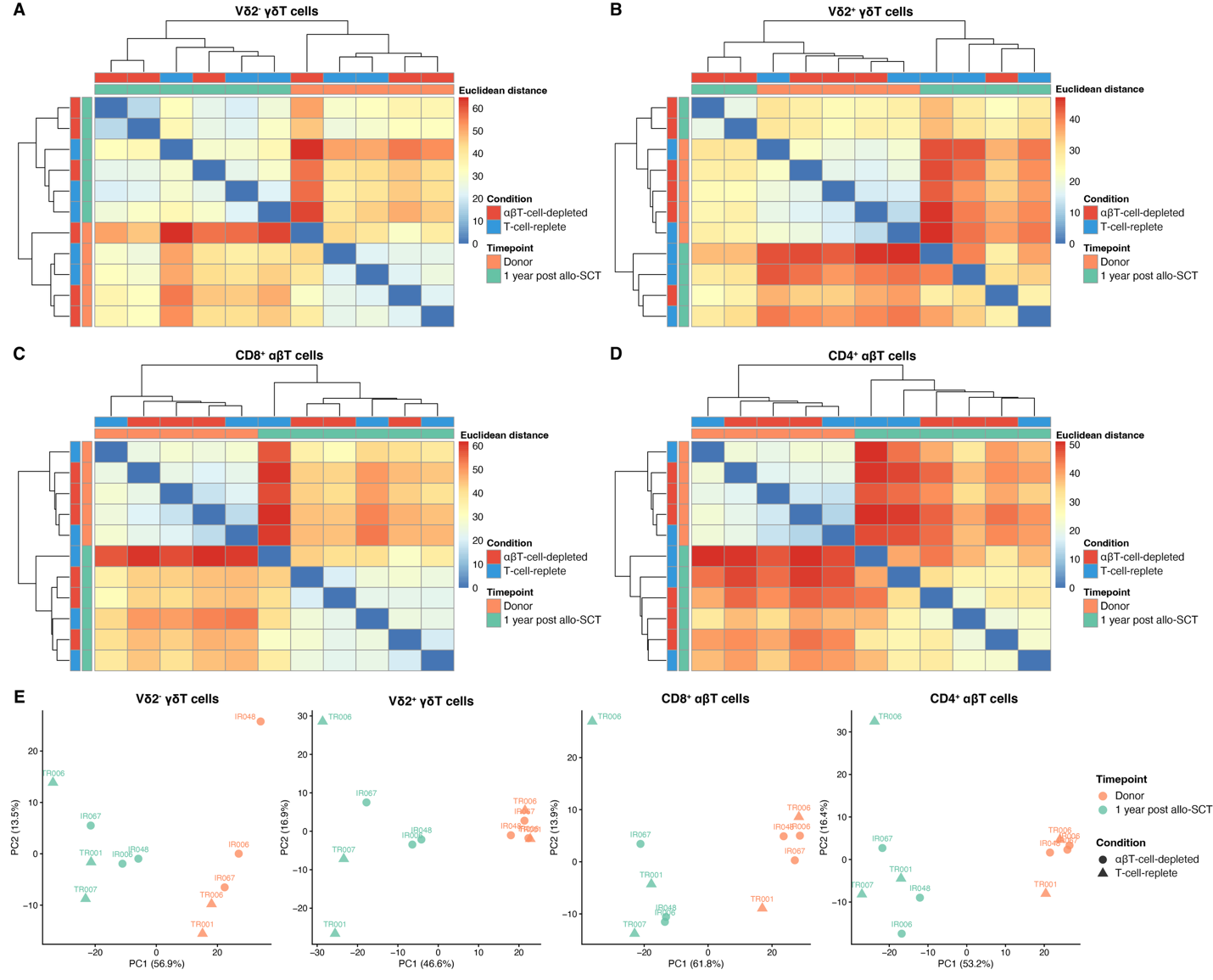
***Supplementary Figure S5. Sample-to-sample Euclidean distance heatmaps and principal component analysis of four T cell subtypes comparing donor baseline and 1 year post allo-HSCT across graft types.*** *Distance calculations and PCA were performed using the top 1,000 most variable genes from pseudobulk-aggregated, normalized log-CPM expression values.* ***(A-D)*** *Euclidean distance heatmaps - Hierarchical clustering based on transcriptome-wide distances between samples for* ***(A)*** *Vδ2- γδT cells,* ***(B)*** *Vδ2+ γδT cells,* ***(C)*** *CD8+ αβT cells, and* ***(D)*** *CD4+ αβT cells. Sample annotations indicate timepoint (orange: donor; teal: 1 year post-SCT) and graft condition (red: αβT-cell-depleted; blue: T-cell-replete). Dark red means the greatest transcriptional distance between samples.* ***(E)*** *Principal component analysis (PCA) plots showing the first two principal components for each T cell subset. Points are colored by time point and shaped by graft condition (circles: αβT-cell-depleted; triangles: T-cell-replete), with patient IDs labeled. The percentage of variance explained by each PC is shown on the axes.*

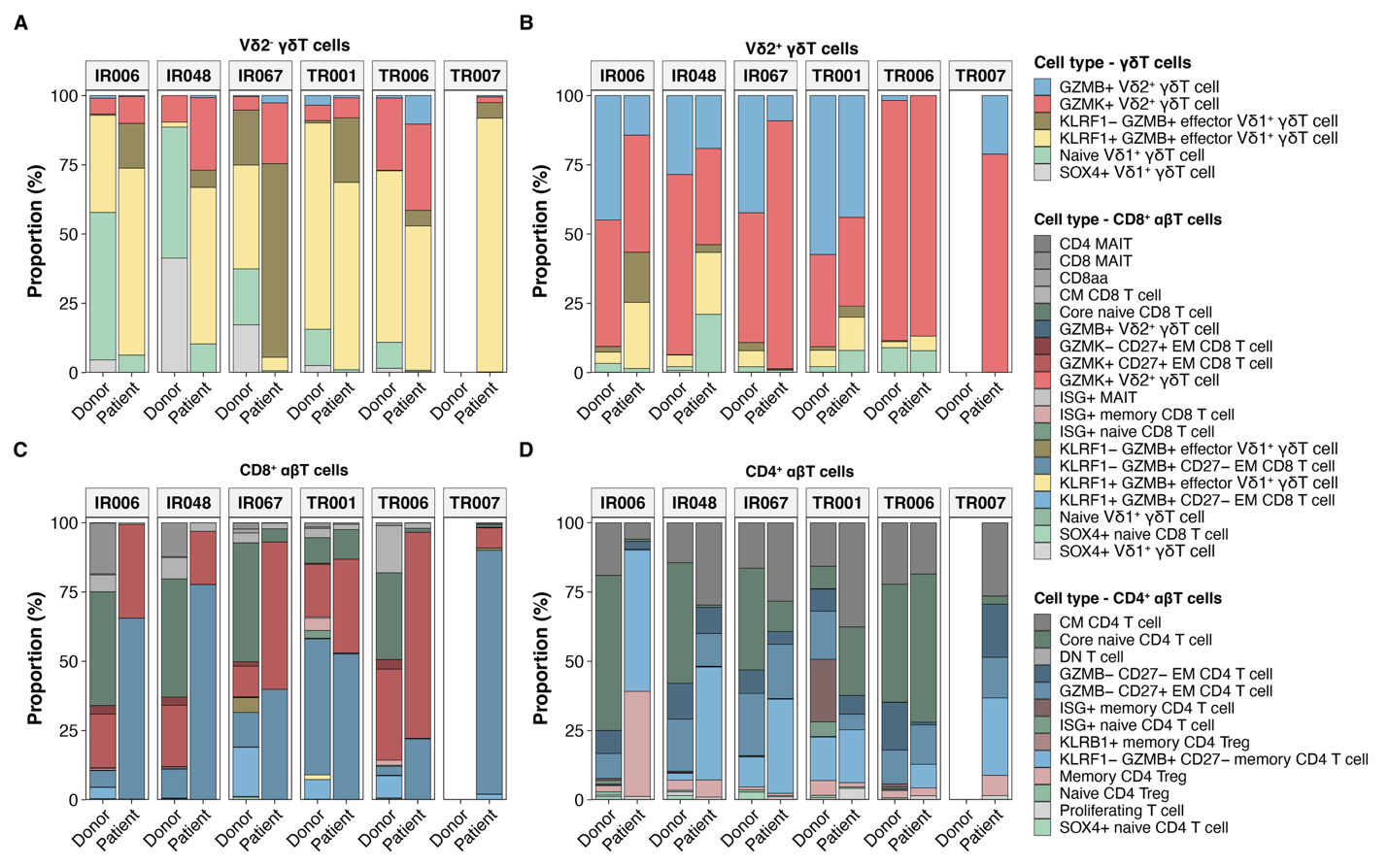

***Supplementary Figure S6. Stacked bar plots showing the proportional distribution of T cell phenotypes in donor grafts and patient peripheral blood at one year post-SCT across six donor-patient pairs.*** *Cell phenotypes were annotated using Azimuth with the Allen Institute Data as a reference dataset.* ***(A)*** *Vδ2- γδT cells,* ***(B)*** *Vδ2+ γδ T cells,* ***(C)*** *CD8+αβ T cells,* ***(D)*** *CD4+ αβT cells. Colors indicate phenotype categories, including naive, memory, effector memory, and cytotoxic (Granzyme+) subsets. Patient TR007 lacks a matched donor sample. Percentages are calculated relative to the total number of cells within each T cell subset per sample.*

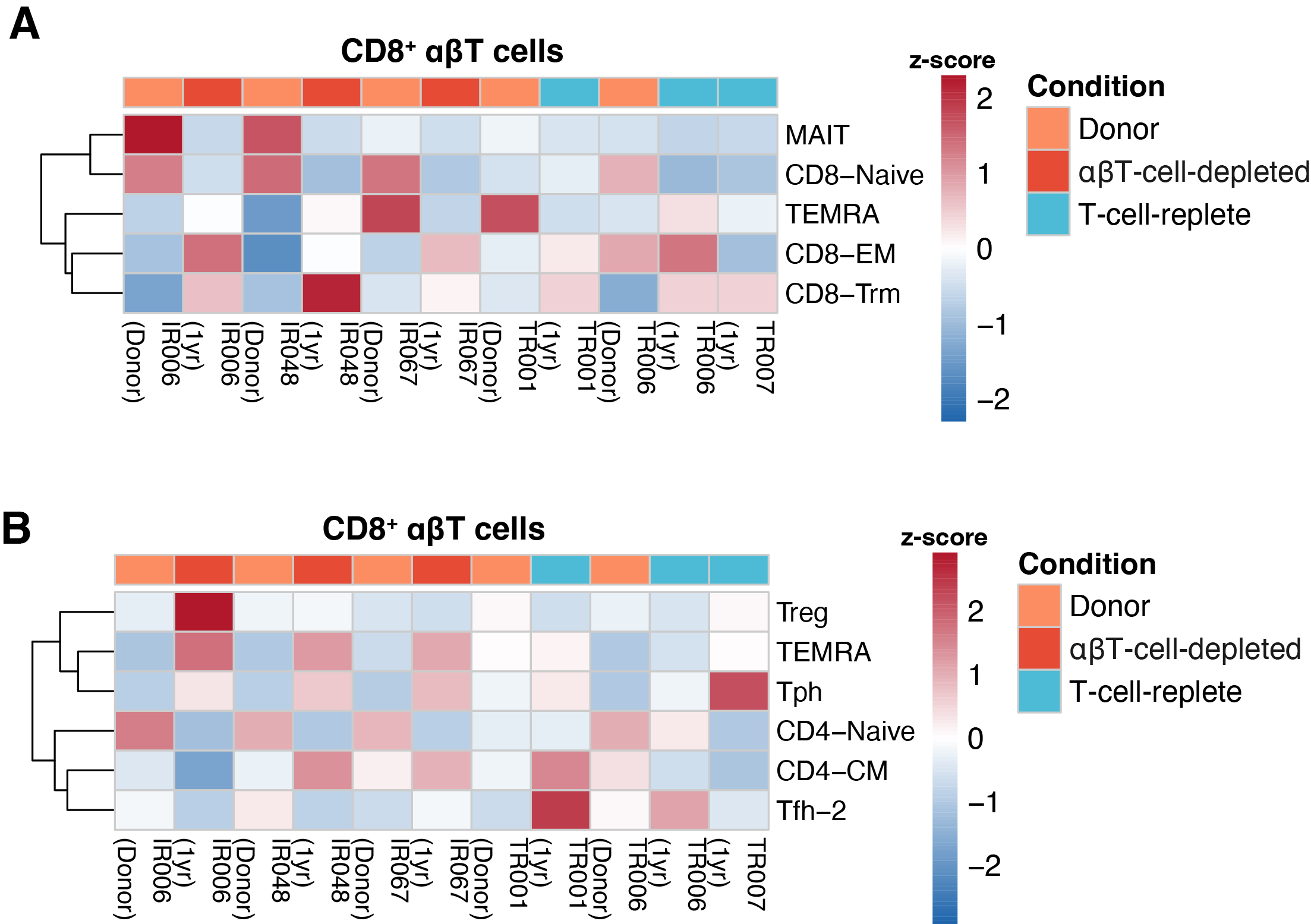

***Supplementary Figure S7. Heatmaps showing average lineage scores*** *for* ***(A)*** *CD8+ αβT cells, and* ***(B)*** *CD4+ αβT cells. Each column represents an individual patient sample at 1 year post-SCT, grouped by transplant type (αβT-cell-depleted: IR006, IR048, IR067; T-cell-replete: TR001, TR006, TR007). Rows represent lineage programmes identified by TCAT. Values are scaled by row (z-score) to highlight relative differences across samples. Color scale: blue (low), white (medium), red (high). Hierarchical clustering of programmes is shown on the left.*

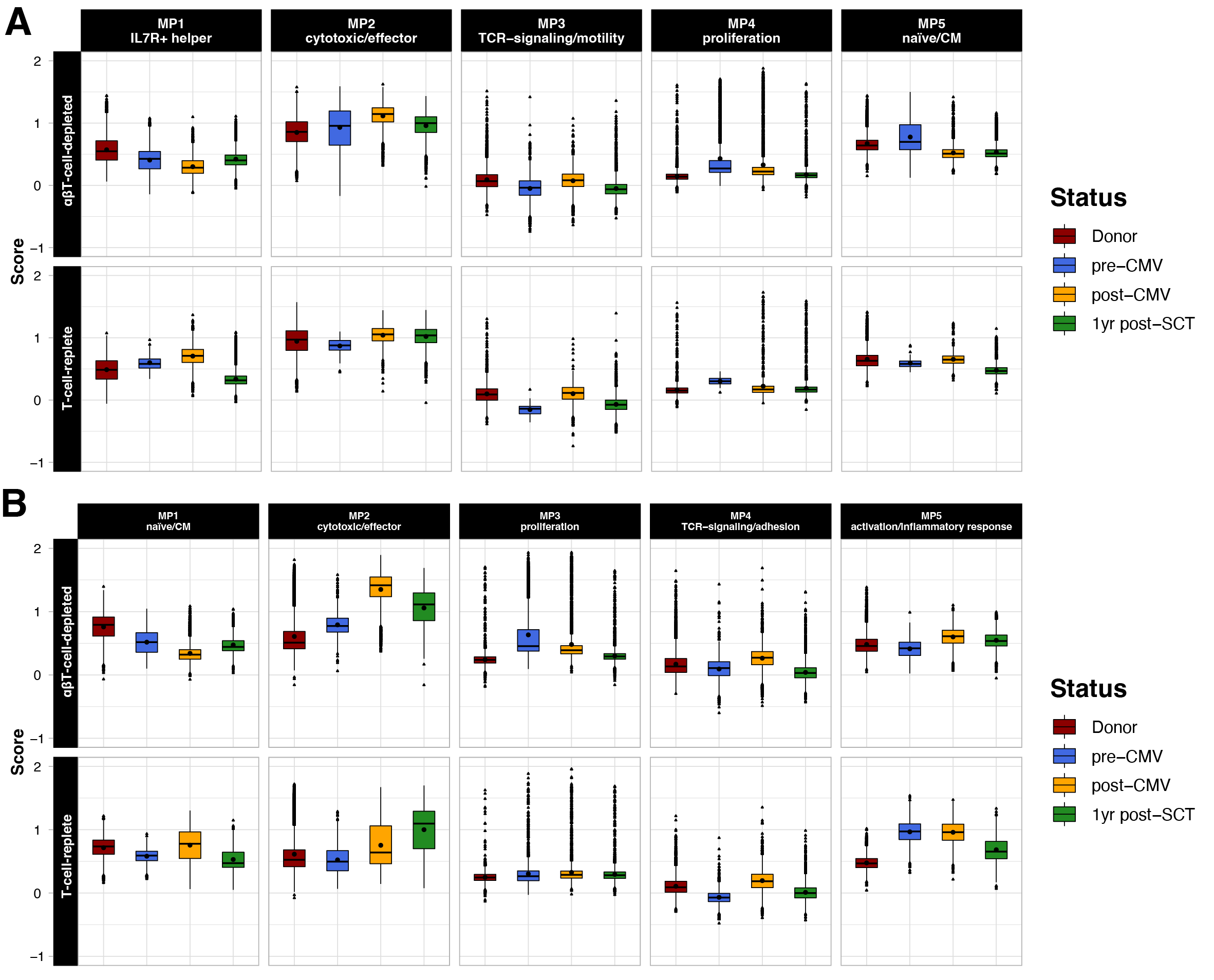

***Supplementary Figure S8. NMF meta-program dynamics during T-cell reconstitution after allo-HSCT.
(A)*** *Single-cell module scores of five γδ T-cell NMF meta-programs (MP1–MP5) are shown as boxplots across donors and post-transplant timepoints, stratified by transplantation platform. MP1 reflects a helper-like γδ program (IL7R, KLRB1, RORA, IL18RAP); MP2 a terminal cytotoxic/NK-like γδ effector program (CCL5, NKG7, PRF1, GNLY, FGFBP2); MP3 a TCR signaling and motility-associated program; MP4 a proliferation/cell-cycle program; and MP5 a naïve/central-memory-like γδ program (TCF7, LEF1, CCR7, BACH2).* ***(B)*** *Single-cell module scores of five αβ T-cell NMF meta-programs (MP1–MP5) are shown as boxplots across donors and post-transplant timepoints, stratified by transplantation platform (stratified by patient in Supplementary Fig. S8). MP1 corresponds to a naïve/central-memory-like program (TCF7, LEF1, IL7R, CCR7); MP2 to a terminal cytotoxic effector program (CCL5, NKG7, PRF1, GNLY, granzymes); MP3 to a proliferation/cell-cycle program (MKI67, PCNA); MP4 to a signaling/adhesion program (FYN, PRKCH, ITGA4, STAT4); and MP5 to an early activation/inflammatory response program (TNFAIP3, ZFP36, DUSP1/2, JUN/JUNB, NR4A2, CD69). Full gene lists for each meta-program are provided in Supplementary Table 3. Colors indicate sampling status. Boxplots show median and interquartile range; points represent individual cells.*

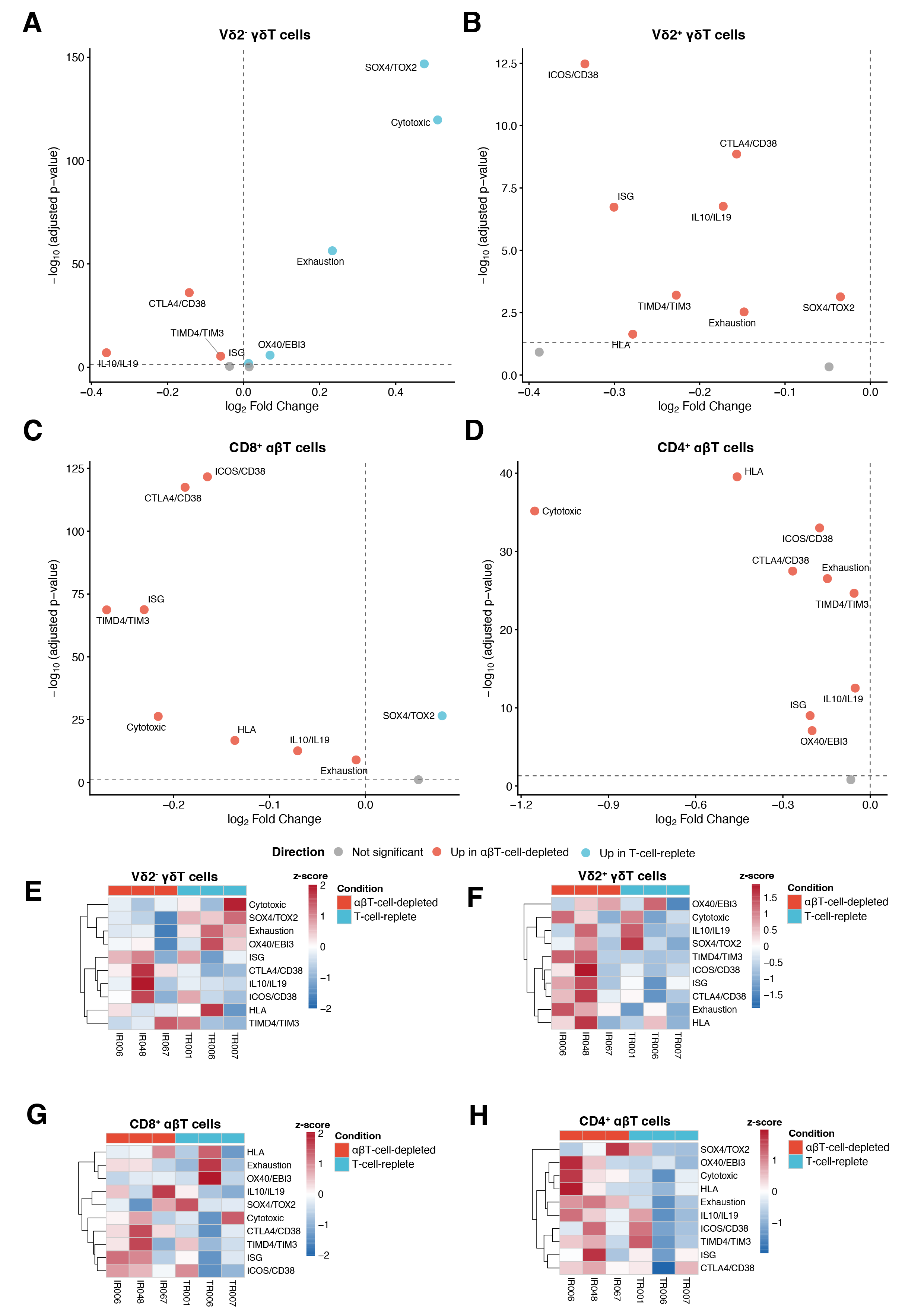

***Supplementary Figure S9. Differential enrichment of functional programmes (A-D)*** *Volcano plots showing differential enrichment of functional programmes between T-cell-replete and αβT-cell-depleted transplants for each T cell subset at 1 year post-SCT. X-axis: log2 fold change (positive values indicate higher in T-cell-replete). Y-axis: -log10(adjusted p-value). Horizontal dashed line indicates FDR < 0.05. Points are colored by direction: blue (higher in T-cell-replete), red (higher in αβT-cell-depleted), grey (not significant). Significantly different programmes (FDR < 0.05) are labeled. Statistical comparison by Wilcoxon rank-sum test with Benjamini-Hochberg correction. Heatmaps showing average functional programme scores for* ***(E)*** *Vδ2- γδT cells,* ***(F)*** *Vδ2+ γδT cells,* ***(G)*** *CD8+ αβT cells, and* ***(H)*** *CD4+ αβT cells. Each column represents an individual patient sample at 1 year post-SCT, grouped by transplant type (αβT-cell-depleted: IR006, IR048, IR067; T-cell-replete: TR001, TR006, TR007). Rows represent individual functional programmes identified by TCAT. Values are scaled by row (z-score) to highlight relative differences across samples. Color scale: blue (low), white (medium), red (high). Hierarchical clustering of programmes is shown on the left.*

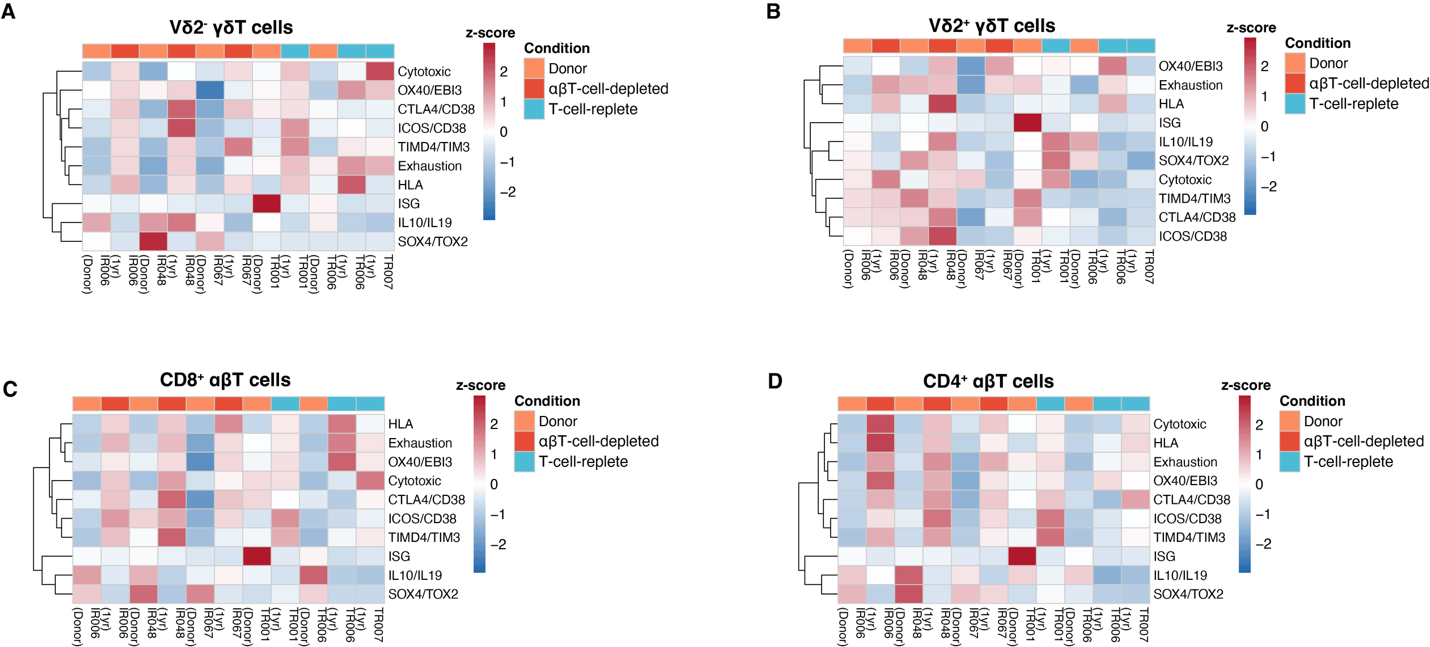

***Supplementary Figure S10.*** *Heatmaps showing average functional programme scores for* ***(A)*** *Vδ2- γδT cells,* ***(B)*** *Vδ2+ γδT cells,* ***(C)*** *CD8+αβ T cells, and* ***(D)*** *CD4+ αβT cells. Each column represents an individual patient sample at 1 year post-SCT, grouped by transplant type (αβT-cell-depleted: IR006, IR048, IR067; T-cell-replete: TR001, TR006, TR007). Donor samples are shown for reference where available. Rows represent individual functional programmes identified by TCAT. Values are scaled by row (z-score) to highlight relative differences across samples. Color scale: blue (low), white (medium), red (high). Hierarchical clustering of programmes is shown on the left.*
